# Neuroendocrine arylhydrocarbon receptor regulates gut microbiome of *C. elegans* via redox tone

**DOI:** 10.64898/2026.06.18.733066

**Authors:** Ciara Hosea, Adrien Assié, Fan Zhang, Buck S. Samuel

**Affiliations:** Program in Development, Disease Models and Therapeutics, 1 Baylor Plaza, Baylor College of Medicine, Houston, TX 77030, USA; Alkek Center for Metagenomics and Microbiome Research and Department of Molecular Virology and Microbiology, Baylor College of Medicine, 1 Baylor Plaza, Houston, TX 77030, USA; Department of Biological Sciences, Louisiana State University, 1146 Pleasant Hall, Baton Rouge, LA 70802

## Abstract

The gut microbiome profoundly influences host health, with disrupted microbial communities implicated in inflammatory, metabolic, and neurological diseases. The aryl hydrocarbon receptor (AHR) shapes intestinal immunity and microbial composition, yet the mechanisms by which AHR mediates selective microbiome assembly remain poorly understood. Whether distant organ systems such as the nervous system actively participate in microbial community regulation is unexplored.

Here we show that *Caenorhabditis elegans* neuronal AHR-1 orchestrates selective gut microbiome assembly through a neuroendocrine cascade that calibrates intestinal redox tone. Using a defined natural microbiome community, we demonstrate that *ahr-1* loss triggers specific microbiome changes, as *Enterobacterales* and *Pseudomonadales* bloom up to 24-fold while *Sphingobacteriales* decline. Adult-specific AHR-1 depletion recapitulates this dysbiosis, separating the role of AHR-1 in neurodevelopment from its regulation of the microbiome. Neuronal AHR-1 regulates expression of the neuropeptide FLP-8 in oxygen-sensing URX neurons, which in turn signals to the intestine to modulate intestinal redox tone. Elevated intestinal redox tone in *ahr-1* mutants favors growth of oxidative stress-tolerant *Enterobacterales* and *Pseudomonadales*, which also exhibit functional and genomic enrichment for stress resistance and AHR ligand biosynthesis pathways. This establishes a precise feedback circuit whereby AHR-1 senses bacterial metabolic output to regulate colonization by ligand-producing taxa. AHR-1 signaling regulates redox tone through downstream regulation of glutathione S-transferase GST-4 and alpha-arrestin ARRD-11. Intestinal *gst-4* knockdowns had the greatest impact on restoring redox tone and microbiome compositional control in the gut. These findings reveal an evolutionarily ancient molecular circuit where AHR serves as a precision metabolic sensor that coordinates whole organismal physiology to maintain host-microbiome homeostasis.

## INTRODUCTION

The gut microbiome profoundly influences host physiology, modulating immunity, neurodevelopment, and metabolism [1–3]. Disrupted microbial communities are associated with inflammatory bowel disease, metabolic disorders, and neurological conditions including Parkinson’s disease, autism spectrum disorder, and schizophrenia, where patients exhibit characteristic compositional alterations with specific bacterial populations showing distinctive abundance patterns [4–6]. Given these far-reaching consequences, hosts deploy an extensive cadre of receptors to recognize a plethora of microbial metabolites and patterns [7,8]. This monitoring is essential for both maintaining a healthy homeostasis with the gut microbiome and alerting the host to the presence of potential pathogens. The aryl hydrocarbon receptor (AHR) has emerged as a central regulator of xenobiotic metabolism, immunity and metabolic homeostasis [9–11]. Mammalian intestinal AHR senses microbial tryptophan-derived metabolites to modulate immune responses and selectively shape microbial communities [12–14]. However, mechanisms underlying AHR-mediated selective microbiome acquisition remain poorly understood, with studies examining only local intestinal AHR function.

*Caenorhabditis elegans* provides an exceptional platform through its AHR ortholog *ahr-1*, which exhibits exclusive neuronal expression with no intestinal localization, unveiling potential for neuroendocrine control of microbial colonization [15,16]. Extensive characterization establishes *ahr-1* as a master regulator of neuronal development controlling cell fate specification and sensory circuits in oxygen-sensing URX neurons, mechanosensory neurons, and GABAergic motor neurons [17,18]. Critically, *C. elegans* AHR-1 responds selectively to bacterial metabolites including *P. aeruginosa* phenazines and tryptophan derivatives rather than classical xenobiotics [19], suggesting ancestral sensory functions for detecting microorganism-derived signals [20]. Recent work implicates *ahr-1* in pathogen defense, lipid metabolism, intestinal barrier integrity, and aging influenced by bacterial diet [21–23]. With 302 mapped neurons and *ahr-1* expression in body cavity neurons contacting pseudocoelomic fluid positioned as environmental sensors detecting circulating microbial metabolites, *C. elegans* enables investigation of how distant neuronal populations orchestrate gut community assembly. While *ahr-1*’s neurodevelopmental and metabolic functions are established [17,23], its role in selective microbiome acquisition remains unexplored.

This investigation establishes that neuronal AHR-1 functions in adulthood to maintain microbiome homeostasis by monitoring microbial metabolites and coordinating responses that selectively shape community structure. Using a defined natural microbiome of *C. elegans* (BIGbiome, 63-members [24]), we demonstrate that *ahr-1* loss triggers characteristic blooms in AHR ligand-producing taxa, with *Enterobacterales* and *Pseudomonadales* expanding 2.5- to 24-fold while *Sphingobacteriales* decline. Temporal-specific protein degradation reveals AHR-1 functions during adulthood in microbiome surveillance that are distinct from its known neurodevelopmental roles. Further, we identify a neuroendocrine molecular cascade whereby neuronal AHR-1 regulates expression of neuropeptide *flp-8* (tachykinins) in URX oxygen-sensing neurons, with 100-fold transcriptional reduction upon *ahr-1* loss. FLP-8 in turn modulates intestinal redox tone through effectors *gst-4* (glutathione S-transferase) and *arrd-11* (arrestin domain-containing protein 2), plus compensatory responses by intestinal transcription factors *aha-1* (ARNT/HIF-1β) and *pqm-1* (Spalt-like 2). Specifically, intestine-specific knockdown of *gst-4* exhibited the greatest rescue of a wildtype-like microbiome composition in *ahr-1* mutants. Mechanistically, we show that disruption of AHR-1 elevates redox tone allowing more stress-tolerant *Enterobacterales* and *Pseudomonadales* to bloom. Similarly, these taxa also exhibited functional and genomic enrichment for stress resistance and AHR ligand biosynthesis pathways. Our studies thus define highly conserved AHR driven molecular pathways in neuroendocrine regulation of microbiome selectivity and homeostasis.

## RESULTS

### AHR-1 shapes composition and diversity of *C. elegans* adult gut microbiome

While mammalian studies have established AHR as a regulator of intestinal health and microbiome composition [10,12,25,26], the molecular mechanisms through which AHR orchestrates selective microbial community assembly remain undefined. We leveraged *C. elegans* as an experimental platform to dissect these mechanisms, capitalizing on several unique advantages: a compact genome enabling precise genetic manipulation, exclusive neuronal expression of the AHR ortholog *ahr-1* that eliminates confounding intestinal cell-autonomous effects, a simple 302-neuron nervous system with complete anatomical maps facilitating circuit-level analysis, and a tractable natural gut microbiome amenable to controlled manipulation [27]. To systematically assess how *ahr-1* influences microbiome composition, we colonized N2 wildtype and *ahr-1(ia03)* loss-of-function mutant animals with BIGbiome, a representative 63-member microbiome of the natural *C. elegans* microbiome that we have developed [24] (**Table in Table S1**). This defined community enables reproducible experimental interrogation of host-microbe interactions while capturing the taxonomic and functional diversity characteristic of native microbiomes. In *C. elegans*, gut microbes initiate colonization in young adulthood and a stable and selective microbiome distinct from the surrounding environment is observed by day 3 of adulthood [24].

Focusing on mature microbiomes of day 3 adults, 16S rRNA gene sequencing revealed that *ahr-1* mutants harbor profoundly altered gut microbiome compositions compared to wildtype animals. Microbiome community structures diverged dramatically from both the input BIGbiome lawn and wildtype-colonized communities (**Fig 1B**). Principal component analysis (PCA) of bacterial operational taxonomic units (OTUs) demonstrated significant separation of *ahr-1* mutant animals in principal component space (p=0.001), suggesting a specific change in microbiome community states rather than passively acquiring microbes (**Fig 1D**). The divergence between wildtype and *ahr-1* mutant microbiomes demonstrates that neuronal *ahr-1* activity drives selective recruitment rather than passive microbial colonization.

**Fig 1.**
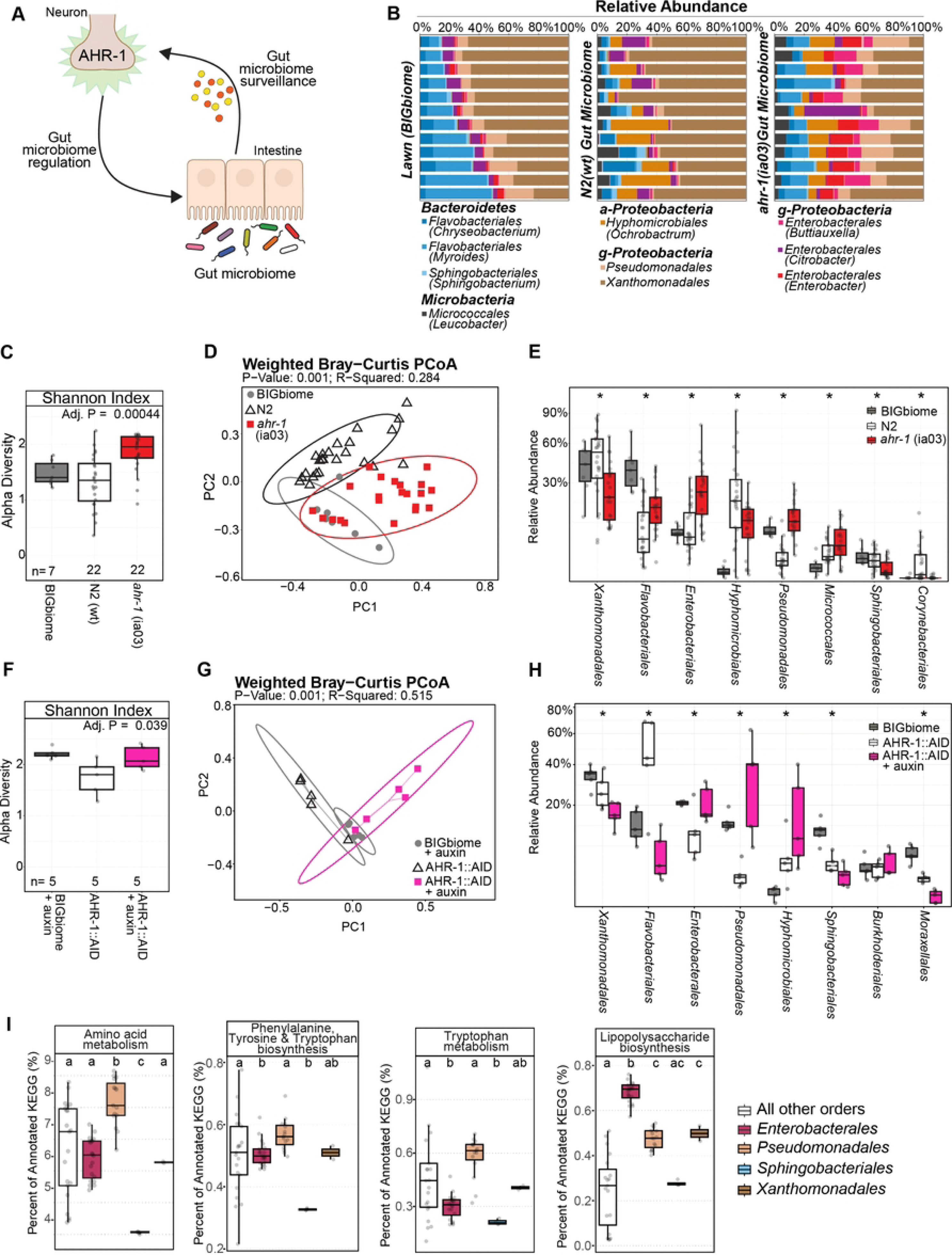
Loss of neuronal AHR signaling causes specific shifts in gut microbiome composition. **(A)** Schematic of interplay between AHR-1 and the gut microbiome. **(B-E)** Previously germ-free animals [N2 (wt) and *ahr-1*(ia03) mutants were raised on BIGbiome lawns and collected as day 3 adults for gut microbiome sequencing. **(F-H)** Post developmental knockdown of AHR-1 was tested using a neuronal Tir1 with an auxin-inducible degron tagged AHR (AHR-1::AID) strain treated with auxin (4mM) from late L4 stage to day 3 of adulthood compared to lawn and untreated controls (no auxin). Microbiome diversity metrics for both sets of strains are noted, including: **(C/F)** within sample alpha diversity (Shannon Index); **(D/G)** PCoA plot of between sample beta diversity metrics (weighted Bray-Curtis dissimilarity); and **(E/H)** relative abundance of bacterial orders. **(I)** Proportion of KEGG pathway genes producing potential AHR ligands in BIGbiome taxa. Kruskal-Wallis test followed by Dunn-Bonferroni post-hoc test for comparison of significance for relative abundance of each order between RNAi conditions represented by compact letter format calculated using microbial order and strain facets. Significance is displayed in the compact letter format. Raw data used to create graphical visualizations can be found in **S1 Data**. 16S sequencing analyses outputs used to create graphical visualizations can be found in **S2 Data**.

Alpha diversity analysis, which quantifies within-sample community richness and evenness, revealed a striking signature of *ahr-1* loss. Mutant animals exhibited significantly elevated Shannon diversity indices (1.86 ± 0.08) compared to wildtype (1.33 ± 0.06, p=0.00044), indicating that *ahr-1* mutants harbor more diverse bacterial assemblages with reduced evenness across constituent strains (**Fig 1C**). This elevated diversity reflects diminished selectivity, where loss of *ahr-1* surveillance permits colonization by a broader spectrum of bacterial taxa that would otherwise be excluded or suppressed. The pattern suggests *ahr-1* normally functions to curate a more restrictive, balanced community composition through active regulatory mechanisms.

Taxonomic profiling at the order level unveiled the specific bacterial populations subject to *ahr-1* regulation. *ahr-1* mutants displayed dramatic and highly significant expansions of three orders: *Enterobacterales*, *Pseudomonadales*, and *Flavobacteriales*, all exhibiting substantial increases in relative abundance (p<0.01) (**Fig 1E**). *Enterobacterales*, which remained relatively constrained in wildtype animals, bloomed to dominate substantial fractions of the *ahr-1* mutant microbiome. Similarly, *Pseudomonadales* underwent marked expansion, achieving fold-changes exceeding 20-fold in some experimental replicates. Conversely, *ahr-1* mutants exhibited significant depletions of *Xanthomonadales* (p<0.05), *Hyphomicrobiales* (p<0.01), *Sphingobacteriales* (p<0.05), and *Corynebacteriales* (p<0.05), revealing that *ahr-1* activity promotes colonization by these orders while suppressing the blooming taxa. These characteristic taxonomic shifts establish *ahr-1* as a critical determinant of selective microbial community assembly, functioning through neuronal surveillance mechanisms to actively curate gut microbiome structure.

### AHR-1 functions in adulthood to regulate microbiome composition

The *C. elegans ahr-1* gene has been extensively characterized for critical roles in neuronal development, including GABAergic motor neuron fate specification, sensory neuron migration, and oxygen-sensing circuit establishment [16–18]. Thus, microbiome alterations in *ahr-1* mutants could result from disrupted neurodevelopment independent of surveillance functions as adults. To distinguish between these distinct roles, we developed AHR-1::AID fusion strains enabling precise temporal control of protein depletion via auxin-inducible degradation. We subjected AHR-1::AID animals to a controlled BIGbiome conditions as before without depletion during normal development to ensure proper neurodevelopment, followed by adulthood-specific AHR-1 depletion achieved through naphthaleneacetic acid (KNAA) exposure from L4 stage until collection at day 3 of adulthood. Control animals received vehicle treatment, maintaining normal AHR-1 levels throughout. This experimental design cleanly separates AHR-1’s potential roles in neuronal circuit assembly from its functions in adult microbial surveillance.

Adulthood-specific AHR-1 depletion phenocopied the microbiome alterations observed in constitutive *ahr-1* mutants, definitively establishing an adult-specific function. Alpha diversity analysis revealed that adulthood AHR-1 knockdown animals exhibited elevated Shannon diversity indices (2.13 ± 0.09) compared to control animals maintaining normal AHR-1 levels (1.64 ± 0.07), mirroring the increased diversity observed in *ahr-1* mutants (**Fig 1F**). Beta diversity analysis via PCoA demonstrated that adulthood knockdown animals clustered independently from controls (p=0.001), confirming distinct community structures despite identical developmental histories (**Fig 1G**). Most strikingly, taxonomic profiling revealed that adulthood AHR-1 depletion recapitulated the precise taxonomic signature of constitutive *ahr-1* mutants: *Enterobacterales* expanded 2.76-fold and *Pseudomonadales* surged 24-fold, while *Xanthomonadales* declined 1.41-fold and *Sphingobacteriales* decreased 1.93-fold (**Fig 1H**). The remarkable concordance between adult-specific knockdown and constitutive mutation phenotypes establishes that AHR-1 actively regulates microbiome composition during adulthood through ongoing surveillance mechanisms rather than through developmental programming of neural circuits. These findings reveal that neuronal *ahr-1* exerts continuous influence over gut microbial community structure throughout the adult lifespan, functioning as a real-time sensor and regulator of microbiome homeostasis.

### Genomic functions of microbes that bloom in AHR mutant gut enriched for potential AHR ligands

Microbial community composition profoundly influences host physiology not merely through taxonomic identity but through the collective metabolic capabilities and bioactive compounds that different assemblages produce. Shifts in community structure can dramatically alter the spectrum of bacterial metabolites, signaling molecules, and immunomodulatory compounds to which the host is exposed, fundamentally changing host-microbe biochemical crosstalk. To investigate the functional consequences of *ahr-1*-dependent microbiome alterations, we performed comparative genomic analysis of the bacterial orders represented in BIGbiome [28], focusing on metabolic pathways capable of producing compounds that might serve as AHR-1 ligands.

To assess potential changes in functional potential of microbiome communities, we quantified the genomic potential for ligand biosynthesis across BIGbiome taxa by analyzing gene copy numbers normalized to genome size, calculating the percentage of each genome dedicated to KEGG-annotated metabolic processes and enzymatic pathways associated with known or predicted AHR ligand production. This approach provides a genome-scale metric of biosynthetic capacity, revealing which bacterial orders are metabolically equipped to produce specific classes of bioactive compounds. Genomic analyses of amino acid metabolism revealed striking differences in biosynthetic capacity across bacterial orders. *Pseudomonadales* devoted significantly higher genomic resources to general amino acid metabolism (8.16% of annotated genes) and aromatic amino acid metabolism (0.57%) compared to all other orders assessed, while *Sphingobacteriales* exhibited significantly lower levels of both (4.05% and 0.33%, respectively) (**Fig 1I**). Tryptophan metabolism, which encompasses potential precursors for AHR ligands like indole, indole-3-acetic acid and tryptamine [29], displayed a similar pattern as well (**Fig 1I**). Finally, we assessed lipopolysaccharide (LPS) biosynthetic capacity, as LPS has been identified as a *C. elegans* AHR-1 agonist [19]. *Enterobacterales* (0.69%), *Pseudomonadales* (0.48%), and *Xanthomonadales* (0.50%) all exhibited substantially higher proportions of LPS biosynthetic genes compared to *Sphingobacteriales* (0.28%) and other orders (0.24%) (**Fig 1I**).

Together, these comparative genomic analyses establish that the bacterial taxa that bloom in *ahr-1* mutants (*Enterobacterales* and *Pseudomonadales*) possess significantly enhanced biosynthetic capacity for producing AHR ligands including tryptophan derivatives, aromatic amino acids, and LPS, while the depleted *Sphingobacteriales* exhibit markedly reduced capacity for these pathways. This metabolic architecture supports a feedback model wherein AHR-1 senses bacterial metabolites to calibrate community composition, with loss of *ahr-1* surveillance permitting expansion of ligand-producing taxa and a loss of microbiome homeostasis (**Fig 1A**).

### Transcriptional profiling of AHR-1 mutants reveals impact on microbial responses and host physiology

To identify the molecular mechanisms linking neuronal *ahr-1* activity to microbiome regulation, we performed genome-wide transcriptional profiling of wildtype and *ahr-1*(ia03) mutant animals reared on either *E. coli* OP50 or BIGbiome community, collected at day 3 of adulthood (**Fig 2A**). This dual-condition design distinguishes core *ahr-1*-regulated genes from microbiome-dependent responses that additionally require *ahr-1* activity.

**Fig 2.**
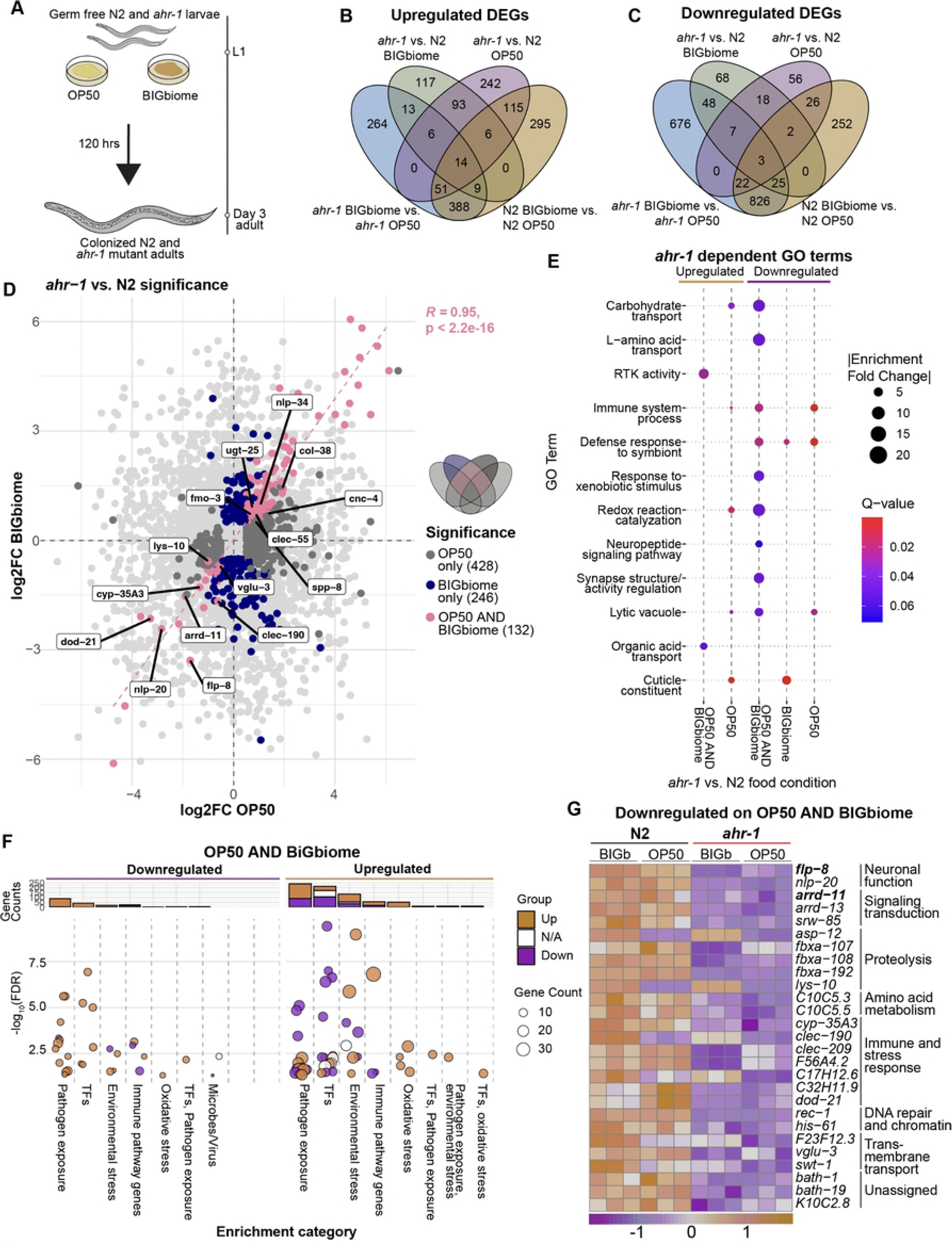
Transcriptional responses to microbes dysregulated in *ahr-1* mutant animals. **(A)** Schematic of experimental setup. Venn diagram of upregulated **(B)** and downregulated **(C)** differentially expressed genes (DEGs) as a function of microbiome and/or host genotype. **(D)** Scatterplot displaying log_2_ fold change (log2FC) differences in *ahr-1* mutant versus N2 (wildtype). DEGs in both microbiome conditions (132, *ahr-1* dependent) are plotted in pink. **(E)** GO enrichment analyses of *ahr-1* dependent DEGs. Color represents Q-value significance, while dot size represents absolute value of enrichment fold change. **(F)** WormExp-based enrichment analyses of existing transcriptional datasets with significant (FDR <0.05) overlap with *ahr-1* dependent DEGs. Barplots detail the count of genes that were upregulated, downregulated, or genetic targets (N/A) in the target dataset. TFs, transcription factors. **(G)** Heatmap displaying *ahr-1* dependent DEGs downregulated genes (27). Key regulators of AHR signaling are noted in bold. Raw data used to create graphical visualizations can be found in **S1 Data**.

From a microbiome response perspective, both genotypes exhibited good concordance between wildtype and *ahr-1* mutant responses on BIGbiome versus OP50 (1,320 genes; R² = 0.95, p < 0.0001; **S2A Fig**). For example, gene ontology analysis identified upregulated genes as being enriched in defense responses to Gram-positive and Gram-negative bacteria, redox reaction catalysis, xenobiotic metabolism, and unfolded protein response regardless of genotype, while cell signaling, carbohydrate transport, and amino acid biosynthesis were downregulated (**S2B Fig**). This suggests that the specific blooms of *Pseudomonas* and *Enterobacteriaceae* in *ahr-1* mutants are reflective of a more selective transcriptional program.

To examine these questions, we focused on genotype-specific analysis of AHR-dependent genes that exhibited coordinate expression patterns across microbial conditions. Specifically, AHR core genes were defined as being differentially expressed in an *ahr-1*-dependent manner in both BIGbiome and OP50 (105 upregulated (**S1A Fig**), 27 downregulated; **Fig 2B–C**, **Fig 2G**) whose fold-changes were tightly correlated (132 genes, R² = 0.95, p < 0.0001; **Fig 2D**). In wildtype animals, BIGbiome upregulated cuticle processes, organic acid transport, and bacterial defense responses. In *ahr-1* mutants, this defensive transcriptional program was largely absent and instead mutant animals exhibit downregulation of genes involved in neuron development, cellular signaling, and neuropeptide signaling (**S2B Fig**).

Enrichment analyses also revealed that *ahr-1* mutants exhibited altered regulation of immune and pathogen response genes. In *ahr-1* mutants on BIGbiome, genes normally induced by *Pseudomonas aeruginosa* PA14 exposure, a pathogen whose defense requires the p38/PMK-1–ATF-7 innate immune axis [30], were significantly downregulated (21 genes, FDR = 2.01×10^-5^, [31]) compared to our OP50 condition. Similarly, genes upregulated in response to

*Enterobacteriales* pathogens like *Serratia marcescens* (176 genes, FDR = 3×10⁻⁴⁴; [32]) and other enteric pathogens like *Enterococcus faecalis* (163 genes, FDR = 3.04×10⁻^21^, [32]) were suppressed in *ahr-1* mutants on BIGbiome as well, compared to OP50 conditions. These downregulated immune-responsive gene sets include known PMK-1 and SEK-1 pathway targets (*pmk-1*, 134 genes, FDR = 2.45×10⁻^67^, [31]; *sek-1*, 321 genes, FDR = 6.62×10⁻⁹⁹, [33]), suggesting that microbiome exposure may stimulate immune responses that are deficient in *ahr-1* mutants (**S2E Fig**). Thus, these microbe responsive genes may help AHR-1 precisely regulate composition of the gut microbiome even though no pathogens are present.

Cross-dataset enrichment analysis revealed striking dysregulation of stress-responsive transcriptional programs in *ahr-1* mutants compared to wildtype on both OP50 and BIGbiome. Among the core AHR genes, we observed significant overlap between genes upregulated in *ahr-1* mutants and genes induced by various oxidative stressors, including hydrogen sulfide exposure (22 genes, FDR=0.00131, [34]), hypoxia (13 genes, FDR=0.0317, [35]), quercetin (9 genes, FDR=0.051, [36]), and tert-butyl hydrogen peroxide (7 genes, FDR=0.00486, [37]) (**Fig 2F**). This is consistent with AHR-1 functioning in oxygen sensing neurons that not only monitor environmental levels of oxygen but also contribute to overall redox balance of the organism.

Lastly, among *ahr-1* downregulated genes (27 genes) specific disruptions in neuroendocrine signaling we also observed. Enriched GO terms included neuropeptide signaling pathway, synapse structure and activity regulation, alongside xenobiotic stimulus response and redox reaction catalysis (**Fig 2E**). Specifically, we identified neuropeptide genes encoding *flp-8* (*FMRFamide*-like peptide; log₂FC = −1.71 OP50, −3.29 BIGbiome; Padj < 5.4×10⁻¹²) and *nlp-20* as being robustly downregulated genes, consistent with prior reports [19,38]. Expression of these neuropeptides overlaps at least partially with AHR-1-positive neurons (i.e., URX; [39]), which could represent a neuroendocrine output of the *ahr-1* transcriptional program regulating gut physiology and microbiome composition.

### Neuronal AHR-1 regulates neuropeptide FLP-8 expression

In addition to transcriptional analyses, multiple lines of evidence support *flp-8* as a transcriptional target of AHR-1. First, *flp-8* and *ahr-1* exhibit overlapping expression patterns in several neuronal populations, most prominently in URX, a bilateral pair of oxygen-sensing neurons critical for aerotaxis and metabolic regulation [40–42] (**Fig 3B**). Second, bioinformatic analysis of the *flp-8* promoter region identified a putative AHR-1:AHA-1 consensus binding site, suggesting direct transcriptional regulation (**Fig 3A**). These convergent observations led us to hypothesize that neuronal AHR-1 coordinates gut microbiome composition through *flp-8*-mediated neuroendocrine signaling.

**Fig 3.**
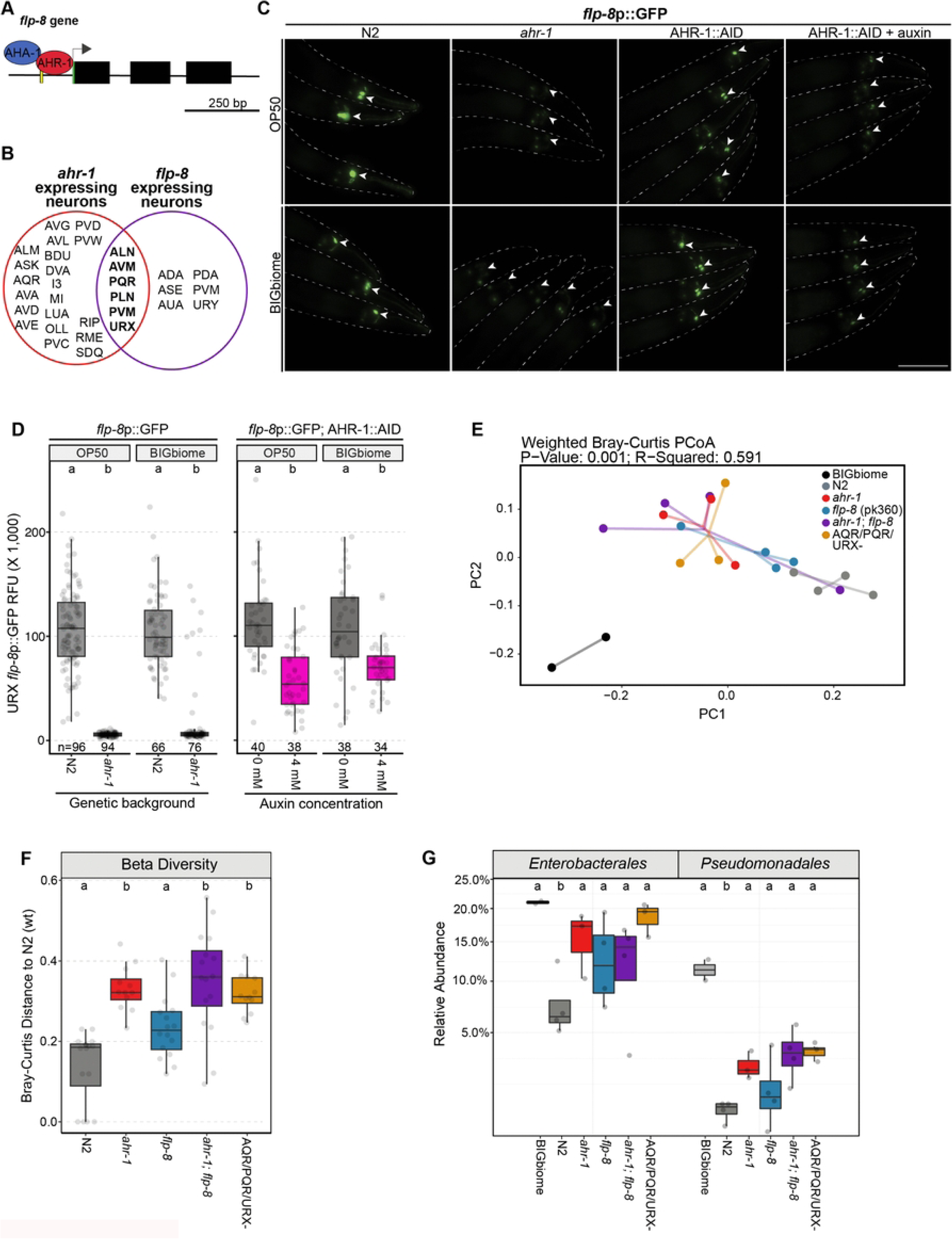
Neuropeptide *flp-8* acts downstream of AHR-1 to regulate the gut microbiome. **(A)** Illustration depicting putative binding site for AHR-1:AHA-1 to the *flp-8* promoter (highlighted in yellow) in neurons, including URX. **(B)** Venn diagram of neuronal expression of *ahr-1* and *flp-8*. **(C-D)** Animals were all raised on *E. coli* OP50 and BIGbiome until day 3 of adulthood. Expression of *flp-8*p::GFP (NY2078) reporter in N2 wildtype, *ahr-1*(ia03) mutant, and AHR-1::AID (+/- 4mM auxin) was assessed by microscopy (scale bar, 100 µm). Arrowheads indicate oxygen-sensing URX head neurons and were quantified in **(D)**. A Kruskal-Wallis test followed by Dunn-Bonferroni post-hoc test was computed for *ahr-1* mutants vs. wt and AHR-1::AID auxin vs. no auxin controls. **(E-G)** Wildtype and mutant animal strains – *ahr-1*, *flp-8* (pk360) mutants, *ahr-1;flp-8* double mutants, and CX7102 animals with genetically ablated AQR, PQR, and URX neurons (AQR/PQR/URX^-^) – were each raised on BIGbiome until day 3 of adulthood. Gut microbiome analyses of animals are displayed for **(E)** PCoA and **(F)** boxplots of beta diversity (weighted Bray-Curtis dissimilarity), plus **(G)** relative abundance of *Enterobacterales* and *Pseudomonadales*. Significance is noted in compact letter display (CLD) format (samples with shared letters = no significant difference); Kruskal-Wallis tests followed by Dunn-Bonferroni post-hoc tests. Raw data used to create graphical visualizations can be found in **S1 Data**. 16S sequencing analyses outputs used to create graphical visualizations can be found in **S2 Data.** Raw flattened images can be found in **S1 Raw Images**.

To validate *flp-8* as an AHR-1 transcriptional target and quantify the regulatory relationship, we generated *flp-8p*::GFP transcriptional reporter animals in both wildtype and *ahr-1* mutant backgrounds. Fluorescence microscopy of URX neurons revealed dramatic dysregulation: *ahr-1* mutants exhibited approximately 100-fold reduction in *flp-8p*::GFP fluorescence compared to wildtype animals on both OP50 and BIGbiome (p<0.0001) (**Fig 3C-D**). This profound transcriptional suppression establishes AHR-1 as a critical positive regulator of *flp-8* expression. However, given *ahr-1*’s well-characterized roles in neuronal development including specification of oxygen-sensing circuits [16], we needed to distinguish whether reduced *flp-8* expression reflects direct adult transcriptional regulation or indirect consequences of developmental circuit defects in URX neurons.

To isolate adult-specific transcriptional effects from potential neurodevelopmental impacts, we deployed the temporal precision of the AID system. We generated *flp-8p*::GFP; AHR-1::AID animals and subjected them to adulthood-specific AHR-1 depletion via auxin exposure as before. Remarkably, adult-specific AHR-1 knockdown phenocopied the *ahr-1* mutants, with auxin-treated animals displaying severely reduced *flp-8p*::GFP fluorescence in URX compared to vehicle-treated controls (p<0.0001) (**Fig 3C-D**). This concordance definitively establishes that AHR-1 actively maintains *flp-8* transcription in adult URX neurons independent of its developmental functions, revealing an ongoing regulatory relationship rather than a developmental programming effect.

Beyond URX, we also observed additional *ahr-1*-dependent modulation of *flp-8* expression in other neuronal populations, with intriguing diet-specific patterns. On BIGbiome, *ahr-1* mutants exhibited significant reductions in *flp-8p*::GFP fluorescence in AUA head interneurons and PVM tail touch neurons (p<0.001), suggesting these neurons also participate in *ahr-1*-coordinated signaling networks. Interestingly, on OP50 monoculture, *ahr-1* mutants displayed the opposite effect in PVM neurons with significantly elevated *flp-8* expression (**S3A-B Fig**), revealing context-dependent and potentially compensatory regulatory mechanisms. Most intriguingly, loss of *ahr-1* triggered ectopic *flp-8* expression in unidentified tail neurons on both microbial diets, a pattern never observed in wildtype animals (**S3C Fig**). To identify these neurons, we crossed *flp-8p*::GFP; *ahr-1* mutant animals into the NeuroPAL background, which provides comprehensive fluorescent labeling of all 302 *C. elegans* neurons with unique spectral signatures enabling computational identification [43]. While NeuroPAL analysis software yielded inconclusive neuronal identification due to the posterior body position and potential spectral overlap, anatomical positioning relative to the rectum suggests these ectopic *flp-8*-expressing neurons likely comprise a combination of PLM mechanosensory neurons, PLN neurons, and/or PQR oxygen-sensing neurons. This ectopic expression pattern suggests that loss of *ahr-1* disrupts normal transcriptional restriction of *flp-8*, potentially representing a compensatory mechanism wherein alternative neuronal populations attempt to restore *flp-8* signaling following its dramatic suppression in URX.

### FLP-8 functions downstream of neuronal AHR-1 regulate microbiome composition

Having established that neuronal AHR-1 directly regulates *flp-8* expression in URX oxygen-sensing neurons, we next tested whether FLP-8 neuropeptide signaling serves as a functional downstream effector of AHR-1’s influence on gut microbiome composition. To dissect the genetic relationship between *ahr-1* and *flp-8* and determine the necessity of *gcy-36*-expressing neurons (AQR/PQR/URX) for microbiome regulation, we performed comprehensive 16S rRNA gene sequencing of five genotypes on BIGbiome at day 3 of adulthood: wildtype animals, *flp-8* single mutants, *ahr-1* single mutants, *ahr-1*;*flp-8* double mutants, and animals with genetically ablated *gcy-36*-expressing neurons (AQR/PQR/URX; [42]). This experimental design enables epistatic analysis to determine whether *flp-8* functions within the *ahr-1* regulatory pathway and whether *gcy-36*-expressing neurons represent the critical cellular locus for neuroendocrine microbiome control.

Beta diversity analysis via principal coordinate analysis demonstrated remarkable concordance among genotypes lacking functional AHR-1 or *gcy-36*-expressing neurons. PCoA ordination revealed that *ahr-1* mutants, *ahr-1*; *flp-8* double mutants, and AQR/PQR/URX^-^ animals clustered tightly together, forming a distinct group that separated clearly from wildtype and *flp-8* mutant communities (PERMANOVA p=0.001) (**Fig 3E**). Quantification of community divergence using Bray-Curtis dissimilarity distances reinforced this pattern: *ahr-1* mutants (distance 0.32), *ahr-1*; *flp-8* double mutants (0.35), and AQR/PQR/URX⁻ animals (0.30) exhibited statistically indistinguishable distances from wildtype, all significantly greater than wildtype self-comparison (0.19) or *flp-8* mutants (0.23), which remained close to wildtype composition (**Fig 3F**). The tight clustering and equivalent divergence of these three genotypes establishes that disruption of the AHR-1 / FLP-8 signaling axis in *gcy-36*-expressing neurons produces a distinct microbiome signature.

Taxonomic profiling revealed that this shared microbiome signature manifests as consistent alterations in specific bacterial orders. All three genotypes with compromised AHR-1/*gcy-36* function (*ahr-1* mutants, *ahr-1*; *flp-8* double mutants, and AQR/PQR/URX^-^ animals) exhibited pronounced expansions of *Enterobacterales*, averaging 2.5-fold above wildtype levels. Most strikingly, these same three genotypes displayed dramatic *Pseudomonadales* blooms substantially exceeding wildtype, while *flp-8* single mutants showed only modest *Pseudomonadales* elevation averaging approximately 3-fold (**Fig 3G**). The remarkable similarity in both the magnitude and specific taxonomic composition of microbiome alterations across *ahr-1* mutants, *ahr-1*;*flp-8* double mutants, and AQR/PQR/URX^-^ animals strongly supports a linear pathway model wherein AHR-1 activity in *gcy-36*-expressing neurons regulates *flp-8* expression, which then functions as a critical downstream neuropeptide effector to shape gut microbial communities.

Critically, the observation that *ahr-1*;*flp-8* double mutants phenocopy *ahr-1* single mutants without enhancement or suppression indicates that *flp-8* functions epistatically downstream of *ahr-1* rather than in an independent pathway. Furthermore, the functional equivalence between genetic *ahr-1* mutation and physical ablation of *gcy-36*-expressing neurons establishes these oxygen-sensing neurons as the essential cellular locus where AHR-1 integrates microbial metabolite sensing with neuroendocrine control of gut microbiome composition. Together, these genetic epistasis experiments define a neuroendocrine regulatory axis where AHR-1 actins in *gcy-36*-expressing neurons to regulate FLP-8 neuropeptide signaling that regulates selective gut microbiome assembly in the intestine.

### AHR-1 and FLP-8 regulate host redox tone in response to microbiome

The URX oxygen-sensing neurons, which co-express *ahr-1* and *flp-8* and serve as the primary locus where AHR-1 regulates *flp-8* transcription, have established roles in neuroendocrine signaling beyond oxygen sensing [41,44]. Recent work has revealed that URX neurons directly detect oxidative stress through peroxide-mediated activation, with URX neuronal activity increasing upon H₂O₂ exposure, positioning these neurons as environmental sensors capable of monitoring redox perturbations [45,46]. Complementing this finding, *ahr-1* has been implicated in both sensing and mitigating reactive oxygen species in *C. elegans* [47]. The convergence of oxidative stress sensing in URX neurons, AHR-1’s documented redox functions, and the dramatic reduction of *flp-8* neuropeptide expression in *ahr-1* mutants led us to hypothesize that the AHR-1 / FLP-8 neuroendocrine axis regulates gut microbiome composition through modulation of intestinal redox tone. Specifically, we reasoned that loss of *ahr-1* might disrupt the host’s ability to sense or respond to microbial-induced oxidative stress, creating intestinal redox environments that selectively favor or disfavor particular bacterial taxa.

To directly measure host redox tone in response to microbial communities, we employed the Grx1-roGFP2 glutathione redox sensor (strain JV2), which provides ratiometric quantification of the GSSG/2GSH ratio, reflecting cellular redox potential and overall redox tone [48]. Grx1-roGFP2 analysis revealed striking microbiome-dependent regulation of host redox tone.

Wildtype animals maintained significantly lower oxidized redox tone on BIGbiome compared to OP50, indicating that complex microbial communities promote a more reduced intestinal environment (Padj=6.86x10^-13^). This microbiome-dependent redox calibration was completely abolished in *ahr-1* mutants (Padj=0.482). On BIGbiome (composed of multiple ligand-producing taxa), *ahr-1* mutants failed to achieve the reduced redox state characteristic of wildtype animals (approximately 1.4-fold increase from OP50, **Fig 4A-B**). Instead, *ahr-1* mutants, *flp-8* mutants, and *ahr-1*; *flp-8* double mutants all exhibited significantly elevated oxidized redox tone (approximately 1.3-fold) compared to wildtype (Padj=0.00125, 0.0303, 7.81x10^-4^, **Fig 4A-B**). On OP50, however, *ahr-1* mutants displayed reduced oxidized redox tone (approximately 2-fold decrease) compared to wildtype, *flp-8* mutants, and *ahr-1*; *flp-8* double mutants (Padj=1.83x10^-4^, 6.43x10^-4^, 0.0129, **Fig 4A-B**). The minimal redox differences across genotypes on OP50 suggests a lack of AHR ligands necessary to robustly engage AHR-1 signaling, rendering the pathway relatively inactive under these conditions.

**Fig 4.**
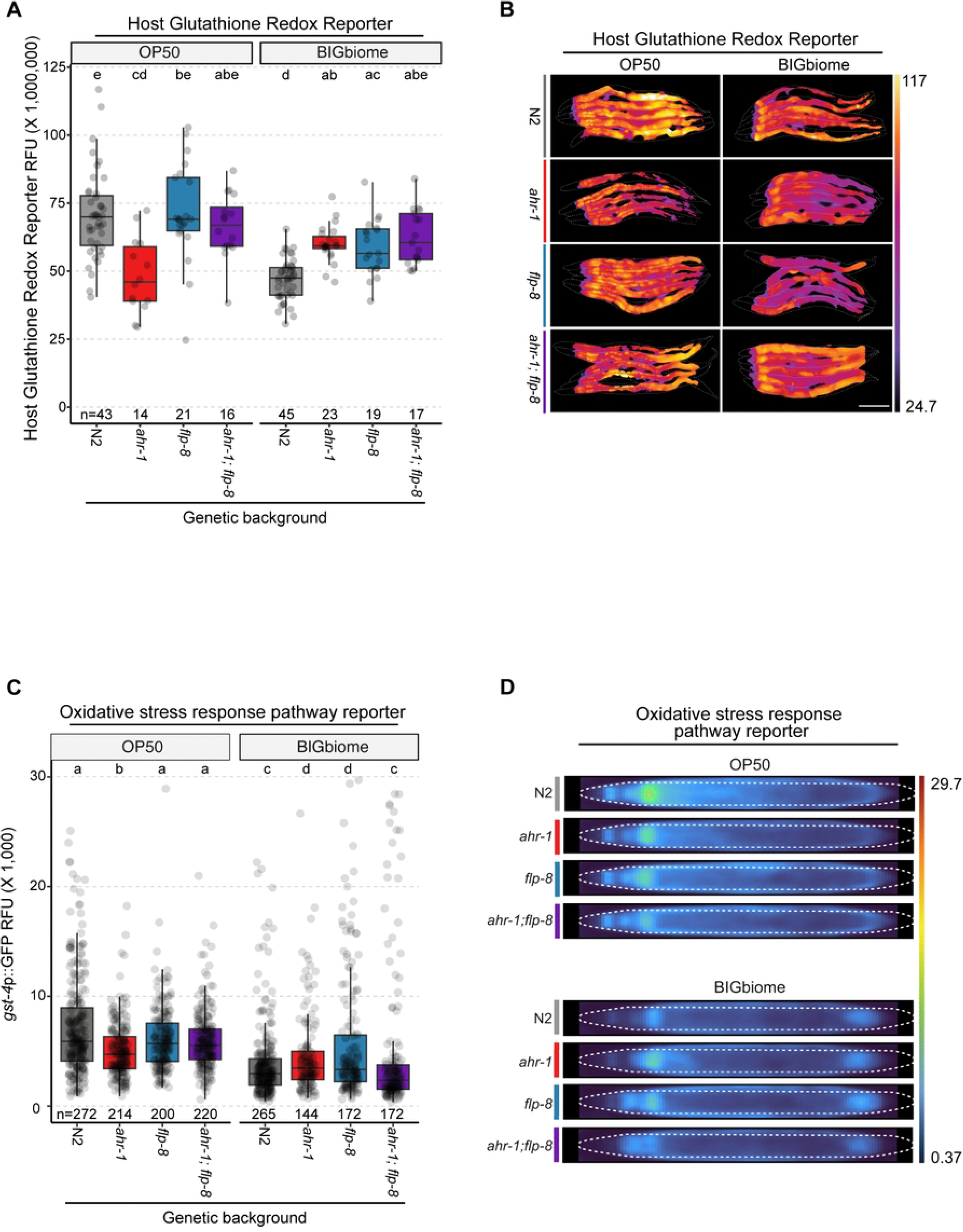
Neuroendocrine AHR signaling regulates redox tone in the intestine. Animals were raised on *E. coli* OP50 or BIGbiome as in Fig 3, plus all strains were crossed with reporters of animal redox tone (JV2, Grx1-roGFP2 glutathione redox sensor) and general oxidative stress (glutathione s-transferase-4, *gst-4*p::GFP). **(A-B)** Fluorescence quantification and representative scaled images of redox tone (oxidization level based on plasma color scale lookup table (LUT)). Scale bar is 250 µm. **(C-D)** Fluorescence quantification and summed scaled images (of all animals in condition) of *gst-4*p::GFP reporter strains. Significance is noted in compact letter display (CLD) format (samples with shared letters = no significant difference); Kruskal-Wallis statistical tests followed by Dunn-Bonferroni post-hoc tests for comparison of fluorescence in different genetic backgrounds for each set of mutants. Raw data used to create graphical visualizations can be found in **S1 Data**. Uncropped flattened images can be found in **S1 Raw Images**.

Next, we assessed redox tone in AHR-1::AID animals with adulthood-specific protein depletion. Strikingly, on the more relevant BIGbiome condition, adulthood AHR-1 depletion completely abolished microbiome-dependent reductions in redox tone, as knockdown animals exhibited approximately 1.3-fold elevation in oxidized redox tone compared to vehicle controls (Padj=0.0208, **S4A-B Fig**). On OP50, adult-specific AHR-1 knockdown produced no significant change in redox tone and were consistently elevated for both OP50 and BIGbiome, suggesting that mutant phenotypes observed for OP50 may be either driven by the neurodevelopmental changes of *ahr-1* mutants or lack of AHR engagement. Together these findings reveal that the AHR-1 / FLP-8 neuroendocrine axis enables hosts to differentially modulate intestinal redox environments in response to ligand-producing microbes. The modulation of intestinal redox tone may therefore be a driver of microbiome community structure.

### AHR-1 regulates oxidative stress responses in a microbiome-dependent manner

To assess how oxidative stress response pathways respond to redox tone alterations, we utilized a *gst-4p*::GFP transcriptional reporter monitoring glutathione S-transferase 4 promoter activation, a canonical antioxidant pathway gene [49]. Previous studies have also implicated *gst-4* in AHR-1 dependent basal responses to reactive oxygen species under OP50 conditions [47]. Overall, analyses of *gst-4p*::GFP expression in wildtype and mutant animals revealed patterns strikingly parallel to redox tone changes, with microbiome-dependent regulation driven both *ahr-1* and/or *flp-8* (**Fig 4C-D**). Though overall low levels of expression for *gst-4* were observed, wildtype animals still did display significantly lower *gst-4* expression on BIGbiome compared to OP50 (approximately 2-fold decrease, Padj=3.13x10^-29^). On OP50, *ahr-1* mutants exhibited significantly reduced *gst-4* activation compared to wildtype (Padj=7.51e-4), which was abolished in *ahr-1*;*flp-8* double mutants (Padj=1, **Fig 4C-D**). On BIGbiome, both *ahr-1* and *flp-8* single mutants exhibited higher *gst-4* expression than wildtype, while *ahr-1*;*flp-8* double mutants exhibited expression similar to wildtype (Padj=0.266). Together, this suggests that AHR-1 regulates GST-4 oxidative stress responses in a microbiome-dependent manner.

Adult-specific AHR-1 depletion revealed notable distinctions from *ahr-1* mutants, with generally elevated *gst-4* expression among the AID strains. On OP50, adulthood knockdown triggered elevated *gst-4* expression (∼1.3 fold, p<0.001), while reduced *gst-4* expression (∼2.3-fold decrease, p<0.001) was observed on BIGbiome. This reduction on BIGbiome does however mirror the oxidized redox tone shift in AHR-1::AID animals on BIGbiome (**S4C-D Fig**). The parallel changes in Grx1-roGFP2 measurements and *gst-4* expression following adult-specific AHR-1 depletion may indicate that GST-4 enzymatic activity helps maintain the reduced redox state in response to BIGbiome engagement of AHR-1.

### Peripheral tissue effectors of AHR-1 neuroendocrine control of redox homeostasis

Having established that neuronal AHR-1 regulates host redox tone through the FLP-8 neuropeptide, we next sought to identify the peripheral molecular machinery that receives and transduces this neuroendocrine signal to modulate redox environments. To identify candidate peripheral effectors, we selected genes based on *ahr-1* dependent differential expression or gene set enrichment in our RNA sequencing datasets (**Fig 2**) with known roles in redox or metabolic regulation. This approach yielded four compelling candidates: *aha-1* (mammalian ortholog ARNT, the AHR-1 heterodimerization partner expressed in multiple peripheral tissues including the intestine [15]); *arrd-11* (mammalian ortholog ARRDC2/TXNIP), an arrestin domain-containing protein downregulated in *ahr-1* mutants (**Fig 2G**)); *skn-1* (mammalian ortholog NRF2, the canonical oxidative stress response transcription factor [50]); and *pqm-1* (related to mammalian SALL2, a transcription factor expressed in intestine that regulates stress responses [51] and the *C. elegans* gut microbiome [24]).

To determine whether these peripheral genes functionally interact with the AHR-1 pathway to regulate redox homeostasis, we performed RNAi knockdowns of each gene or vector controls (L4440) outside the nervous system in AHR-1::AID animals carrying either Grx1-roGFP2 or *gst-4p*::GFP reporters. Animals experienced RNAi during development (L1 to L4 larval stages), then were transferred to BIGbiome with or without auxin to induce adult-specific AHR-1 depletion, and collected at day 3 of adulthood (refer to methods). For redox tone (Grx1-roGFP2), all RNAi conditions displayed significantly increased oxidized redox tone upon AHR-1 depletion compared to their respective no-auxin controls (**Fig 5A-B**). Interestingly, peripheral knockdown of *aha-1*/ARNT significantly elevated redox tone (∼1.25-fold increase, Padj<2.67x10^-3^) compared to empty vector controls both with and without adult-specific AHR-1 depletion (**Fig 5A-B**). This suggests that *aha-1/ARNT* may function independently in the intestine to maintain reduced redox tone regardless of neuronal AHR-1 status. Knockdown of transcription factors known to be involved in oxidative stress responses, *skn-1*/NRF2 and *pqm-1*/SALL2 had no impact on redox tone in any condition tested (**Fig 5A-B**), which could reflect the relatively mild elevations in redox state are not enough to engage these pathways.

**Fig 5.**
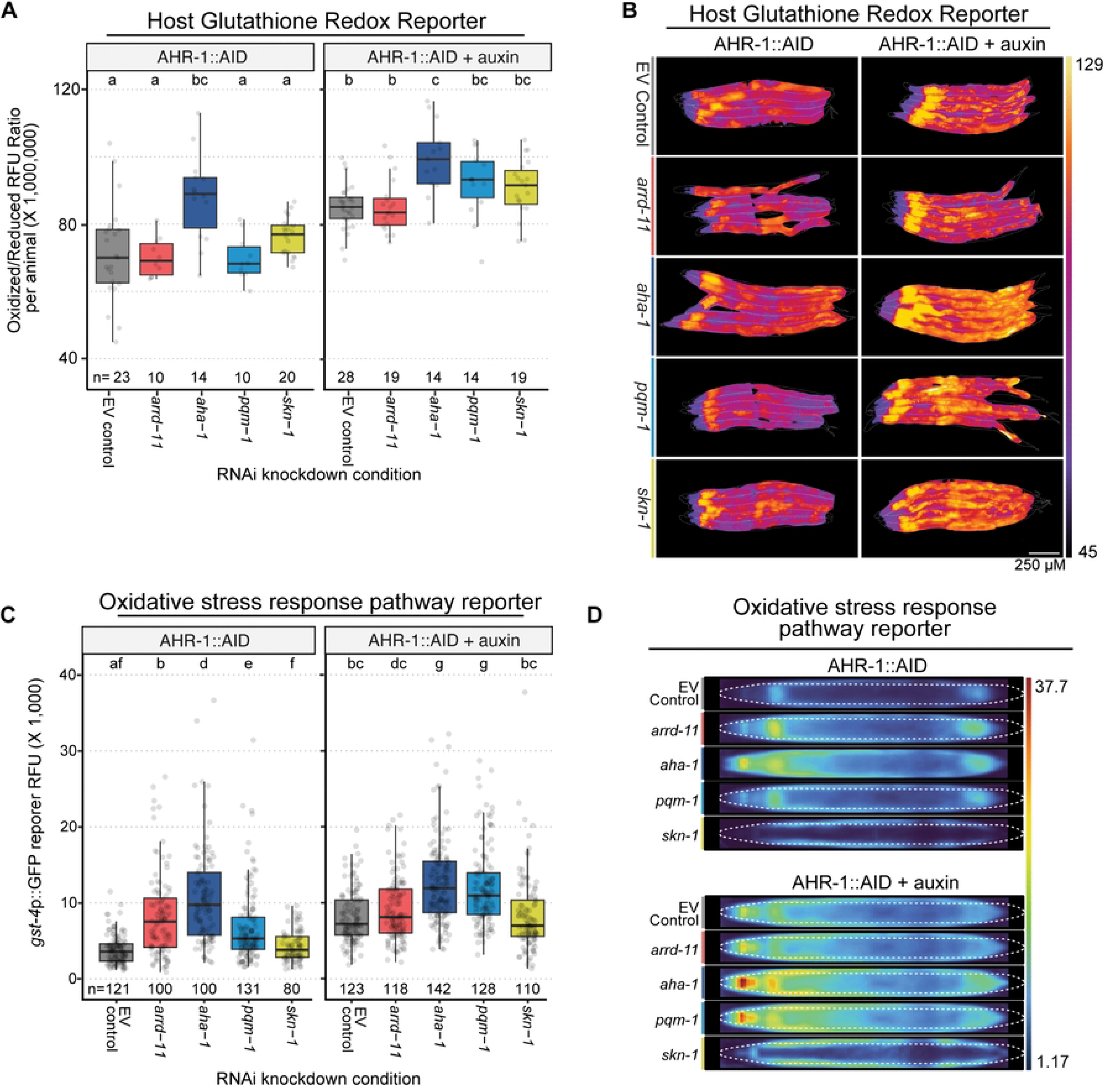
Interplay of downstream mediators on AHR regulation of redox tone in the intestine. Whole animal RNAi knockdowns in AHR-1::AID animals with redox reporters were completed using dsRNA expressing in *E. coli* HT115 during development including empty vector controls (EV, L4440), transferred to BIGbiome at L4 stage (+/- 4mM auxin) and analyzed as day 3 adults as in Fig. 4. **(A-B)** Fluorescence quantification and representative scaled images of redox tone are (AHR-1::AID; Grx1-roGFP2; oxidization level based on plasma color scale Look-up Table (LUT)). Scale bar is 250 µm. **(C-D)** Fluorescence quantification and summed scaled images (of all animals in condition) of AHR-1::AID;*gst-4*p::GFP reporter strains. Significance is noted in compact letter display (CLD) format (samples with shared letters = no significant difference); Kruskal-Wallis tests followed by Dunn-Bonferroni post-hoc test for comparison of fluorescence in different RNAi knockdowns. All pairwise analyses of each auxin-treated condition (AHR-1 depleted) and their respective untreated (no auxin) controls demonstrated significant differences, with the exception of *aha-1*. Raw data used to create graphical visualizations can be found in **S1 Data**. Uncropped flattened images can be found in **S1 Raw Images**.

Next we assessed the impact of these genes on *gst-4p*::GFP transcriptional reporter expression. In wildtype animals, peripheral knockdowns of *arrd-11*/ARRDC2, *aha-1*/ARNT, and *pqm-1*/SALL2 all produced significantly elevated *gst-4::GFP* expression compared to empty vector controls and similar to that observed in AHR-1 depleted animals (**Fig 5C-D**). Following adult-specific neuronal AHR-1 depletion, both *aha-1*/ARNT and *pqm-1*/SALL2 knockdowns were able to further activate *gst-4::GFP* expression above empty vector controls (Padj=2.49x10^-7^, 1.52x10^-8^), suggesting that both transcription factors are acting in parallel to AHR signaling.

Knockdown of *skn-1*/NRF2, a known regulator of *gst-4*, did not significantly alter *gst-4*::GFP expression under any of the conditions tested and were comparable to controls (**Fig 5C-D**). This is consistent with independent roles of AHR-1 and SKN-1 in regulation of *gst-4* expression under baseline conditions [47]. Notably, knockdown of *arrd-11*/ARRDC2 had no enhancing effect on *gst-4::GFP* expression, indicating that it is a primary downstream effector of AHR signaling.

### Knockdown of intestinal effectors of AHR-1 rescue microbiome dysbiosis

Having established that context dependent impacts of *aha-1*/ARNT, *arrd-11*/ARRDC2, *pqm-1*/SALL2 and *gst-4/*GST on redox tone, we next tested whether these genes function as intestinal effectors mediating neuronal AHR-1 control of gut microbiome composition. We performed intestine-specific RNAi in wildtype (MGH171) and *ahr-1* mutant (BIG0108) backgrounds, with developmental RNAi from L1 through L4 followed by BIGbiome colonization through day 3 of adulthood for 16S rRNA sequencing. First, we validated that that intestine-specific RNAi strains carrying *ahr-1* mutation and fed empty vector control exhibited distinct gut microbiome communities from wildtype controls. Indeed, these strains exhibited a distinct microbiome composition as quantified by Bray-Curtis beta diversity and PCoA clustering (**Fig 6B-C**). This dysbiosis is driven by characteristic blooms of *Enterobacterales* and *Pseudomonadales* (**Fig 6E-F**) coupled with elevated Shannon diversity indices indicating reduced microbiome selectivity (**Fig 6D**). These findings confirm that intestinal RNAi methodology itself does not disrupt the core microbiome phenotypes associated with neuronal *ahr-1* loss.

**Fig 6.**
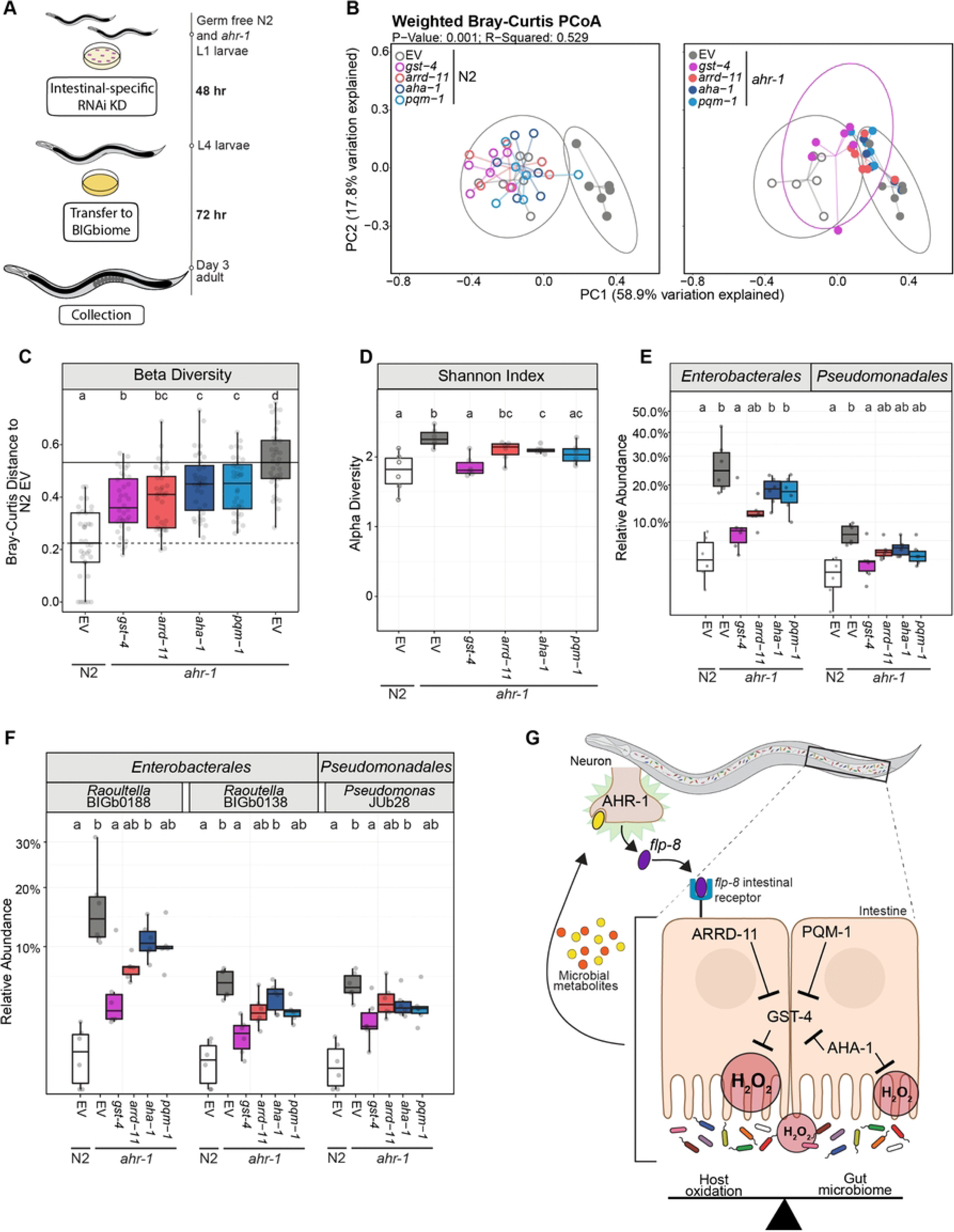
Gut microbiome changes in AHR mutants driven by dysregulation of *gst-4* and *arrd-11*. **(A)** Schematic of experimental setup (similar to Fig. 5) for intestinal-specific RNAi in N2 (wildtype) and *ahr-1(ia03)* mutants. **(B)** PCoA plots of between sample beta diversity metrics (weighted Bray-Curtis dissimilarity) faceted by host genetic background. Box plots of RNAi knockdowns in *ahr-1* mutants that contribute to normalization of gut microbiome metrics toward N2 empty vector (EV, L4440) controls, including: **(C)** weighted Bray-Curtis distances, **(D)** within sample alpha diversity (Shannon Index), and relative abundance of gut bacteria orders **(E)** and strains **(F)** that bloom in ahr-1 mutants. separated by order of bacteria and **(F)** individual species of bacteria of interest. Impact of RNAi knockdowns on N2 strains is displayed in **Fig S5**. Significance is noted in compact letter display (CLD) format (samples with shared letters = no significant difference); Kruskal-Wallis test followed by Dunn-Bonferroni post-hoc test between RNAi conditions. Significance is displayed in the compact letter format. **(G)** Graphical summary highlighting role of AHR-1-dependent regulators ARRD-11 and GST-4, plus supporting responses by PQM-1 and AHA-1 in certain cases. Raw data used to create graphical visualizations can be found in **S1 Data**. 16S sequencing analyses outputs used to create graphical visualizations can be found in **S2 Data.**

Next, we examined the impact of intestinal knockdowns of the candidate AHR-1 effectors. Similar to assays of redox tone, intestinal knockdowns of *aha-1*, *arrd-11*, *gst-4*, and *pqm-1* all had minimal impact on microbiome communities in a wildtype context (**S5A-D Fig**). However, intestinal knockdown of both *gst-4* and *arrd-11* in *ahr-1* mutants significantly rescued most if not all of the wildtype-like microbiome diversity metrics (**Fig 6**). Beta diversity analysis showed *gst-4* knockdown shifted *ahr-1* mutant communities the closest to wildtype controls, while *arrd-11*, *aha-1* and *pqm-1* also had more minor restorations of microbiome structure (**Fig 6B-C**). Alpha diversity measurements revealed *gst-4* and *pqm-1* knockdowns reduced Shannon Indices of microbial richness to wildtype control levels (**Fig 6D**). Taxonomic profiling confirmed that intestinal *gst-4* knockdown in *ahr-1* mutants also suppressed blooms of *Enterobacterales* and *Pseudomonadales* down to the strain level (**Fig 6E-F**), while *arrd-11* knockdowns rescued only *Enterobacterales* levels. Together, the ability of *gst-4* and *arrd-11* to rescue most of the microbiome dysbiosis observed in *ahr-1* mutants positions these redox regulatory genes as key intestinal mediators linking neuronal signals to selective microbiome assembly. More modest but still significant impacts were also generally observed for intestinal knockdown of *pqm-1*/SALL2 and *aha-1*/ARNT in *ahr-1* mutant backgrounds, as both genes showed partial rescue to wildtype beta diversity levels (**Fig 6B-C**) and alpha diversity measures of selectivity (**Fig 6D**). Notably, *pqm-1*/SALL2 knockdown suppressed *Pseudomonadales* blooms in *ahr-1* mutants (**Fig 6E-F**).

Together, these intestine-specific RNAi experiments establish that neuronal AHR-1 remotely controls gut microbiome composition through intestinal effector genes, with *gst-4* functioning as the dominant downstream mediator. Intestinal *gst-4* loss rescues *ahr-1* mutant dysbiosis, which positions GST-4 regulation of redox tone as the main effector through which the AHR-1/FLP-8 neuroendocrine axis shapes microbiome composition.

### Bacterial oxidative stress tolerance and genomic enrichment of oxidative response genes correlates with colonization in *ahr-1* mutants

KEGG Orthology analysis across BIGbiome genomes revealed functional clustering mirroring colonization phenotypes in *ahr-1* mutants. UMAP ordination showed that oxidative stress-resistant *Enterobacterales* and *Pseudomonadales* that bloom in *ahr-1* mutant guts cluster together, while oxidatively sensitive *Sphingobacteriales* that are depleted formed a distinct functional cluster (**Fig 7A**). Differential analyses identified 2,220 features distinguishing these groups (**S1 Data**). *Enterobacterales* and *Pseudomonadales* exhibited elevated proportions of genes for polyketide biosynthesis (log₁₀FDR 1.20), xenobiotic metabolism and glutathione metabolism, plus genes that produce potential AHR ligands like tryptophan metabolism and lipopolysaccharide biosynthesis (**Fig 7B**). Lower colonization of *Sphingobacteriales* in also significantly depleted sphingolipid metabolism genes in the gut microbiome of *ahr-1* mutants (**Fig 7B**).

**Fig 7:**
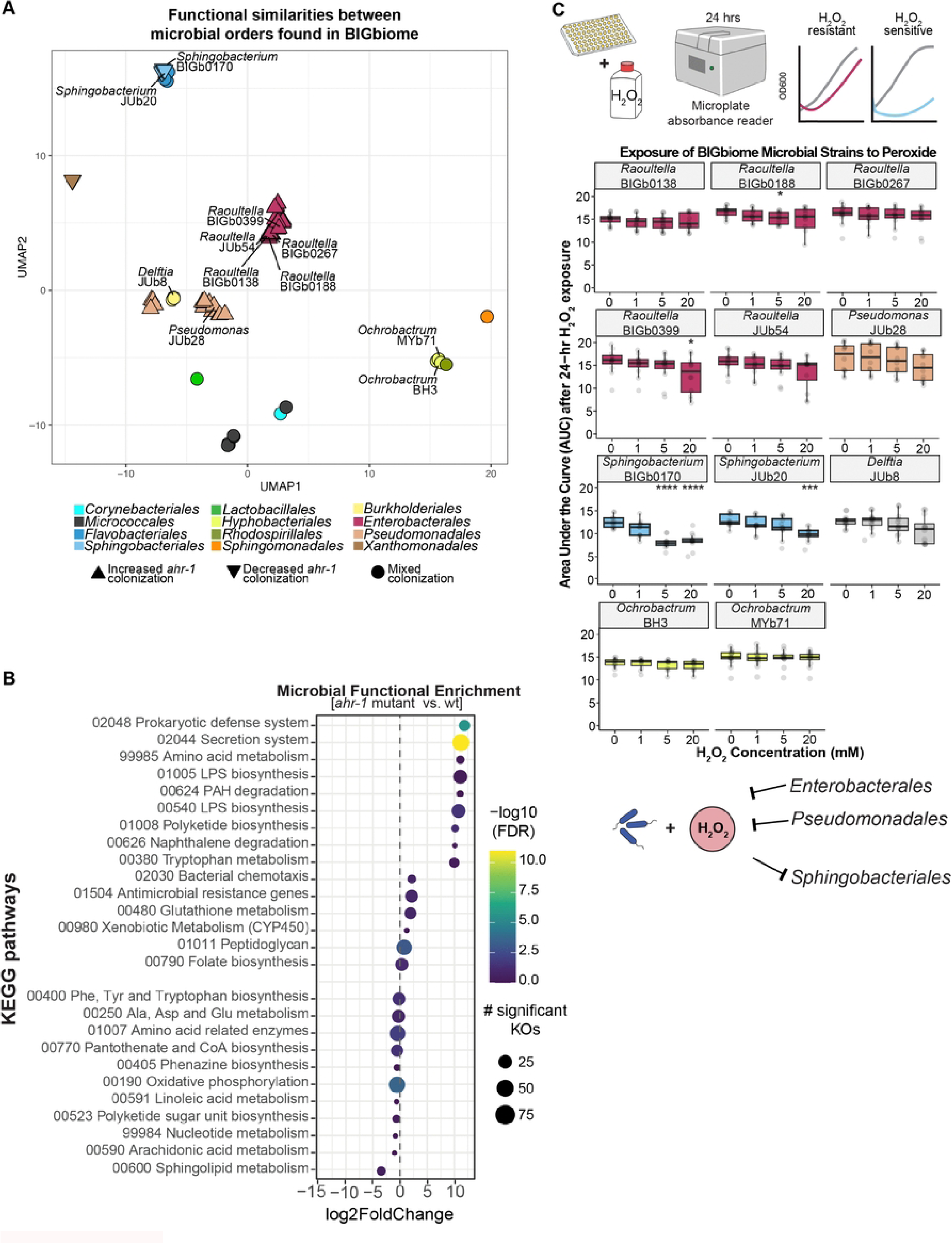
Microbiome strains altered in AHR mutants exhibit genomic enrichment for and enhanced resistance to oxidative stress. **(A)** UMAP plot of genomic clustering of BIGbiome strains based on encoded functions. **(B)** KEGG Ontology (KO) enrichment analyses of genes overrepresented and underrepresented in microbial orders that consistently colonize the *ahr-1* mutant or AHR-1-depleted gut at higher levels (30 strains; *Enterobacterales* and *Pseudomonadales*) versus lower levels (5 strains; e.g., *Sphingobacteriales, Xanthomonadales*). **(C)** Schematic of *in vitro* hydrogen peroxide (H_2_O_2_) resistance assay for microbiome strains. Boxplots display cumulative area under the curve (AUC) values for O.D.600 measurements over 24-hours of exposure to 3% H_2_O_2_ (0mM - 20mM). Statistical significance compared to untreated controls for each strain is noted (Wilcoxon test comparing to control of 0 mM H_2_O_2_; *p<0.05, ***p<0.001, ****p<0.0001). All replicated either 11 or 12 times. Some illustrations created with BioRender.com. Raw data used to create graphical visualizations can be found in **S1 Data**.

To further understand the genomic basis for differential stress tolerance, we focused specifically on bacterial antioxidant and oxidative stress response gene repertoires. Notably, *Enterobacterales* and *Pseudomonadales* displayed elevated counts of glutathione reductase (K00383), glutathione S-transferase (K00799), alkyl-hydroperoxide reductase (K03387), and thioredoxin II (K03672) (Mann-Whitney adjusted p<0.05) compared to other taxa (**S6 Fig**). The enrichment of glutathione-based antioxidant systems in blooming taxa is particularly striking given that intestinal *gst-4* loss rescues *ahr-1* mutant microbiome composition, suggesting a coordinated host-microbe redox axis wherein both host and bacterial glutathione systems establish oxidative environments that selectively favor specific taxa.

To directly test whether differential oxidative stress tolerance explains colonization patterns, we exposed representative bacterial strains to hydrogen peroxide gradients. *Enterobacterales* (*Raoultella* sp. BIGb0138, BIGb0267, BIGb0399), *Pseudomonadales* (*Pseudomonas sp*. JUb28), *Sphingobacteriales* (*Sphingobacterim sp.* BIGb0170 and JUb20), *Hyphomicrobiales* (*Ochrobactrum sp*. BH3 and MYb71), and *Burkholderiales* (*Delftia* JUb8) strains were cultured with 0 mM –20 mM H₂O₂ for 24 hours to determine impact on bacterial growth (**Fig 7C**).

*Sphingobacteriales* strains exhibited the most sensitivity to oxidative stress with BIGb0170 exhibiting the growth inhibition at as little as 2.5 mM H₂O₂ (p<0.05 vs. 0 mM) and JUb20 at 10 mM and greater H₂O₂. In contrast, *Enterobacterales* strains demonstrated robust oxidative stress resistance with no significant growth inhibition across 1–5 mM H₂O₂. Even the most sensitive *Enterobacterales* strain BIGb0399 maintained robust growth until 20 mM H₂O₂. These functional tolerance differences directly correlated with colonization outcomes, as oxidative stress-resistant bacteria like *Enterobacterales* and *Pseudomonadales* bloom in *ahr-1* mutants, while oxidatively sensitive taxa like *Sphingobacteriales* decline.

Together, these functional assays and genomic analyses establish a clear mechanism whereby neuronal AHR-1 regulates intestinal redox tone through the AHR-1/FLP-8/GST-4 axis, creating a selective niche based on bacterial oxidative stress tolerances. The enrichment of glutathione-based antioxidant systems in blooming taxa demonstrates that elevated intestinal oxidation in *ahr-1* mutants specifically favors bacteria equipped to manage peroxide stress through glutathione metabolism, directly linking host neuroendocrine control to functional bacterial trait selection in microbiome assembly.

## DISCUSSION

In this study, we demonstrate that neuronal AHR-1 controls gut microbiome composition in adult *C. elegans* through a neuroendocrine feedback circuit. In the absence of AHR-1, bacteria that are enriched in genomic pathways that produce potential AHR ligands (like *Enterobacterales* and *Pseudomonadales*) bloom in the gut microbiome. Adult-specific depletion of AHR-1 yielded similar microbiome phenotypes that are separable from the role of AHR-1 in neurodevelopment [17]. Following recognition of these bacteria by AHR-1, we identify that AHR-1 activity is centralized in URX sensory neurons where it integrates microbial ligands and coordinates physiologic responses in the intestine by regulation of neuropeptide *flp-8* gene expression. Loss of this AHR-1/FLP-8 neuroendocrine circuit leads to dysregulation of redox tone to a more oxidized baseline, which is driven by changes in intestinal expression of *gst-4/*GSTA and *arrd-11*/ARRDC2. Knockdowns of intestinal *gst-4* and other key factors specifically rescue microbiome dysbiosis phenotypes, identifying GST-4 as the effector bridging neuronal signals to regulation of microbiome homeostasis. Microbial genomic enrichment for oxidative stress genes and heightened resistance to oxidative stress indicates that AHR-1 regulation of redox tone is a driver of microbiome composition. Thus, we establish a distinct brain-gut molecular feedback loop that provides *C. elegans* hosts with specific control over the gut microbiome based on functional properties of its members.

Though less is known for the microbiome, neuronal regulation of intestinal immunity against pathogens is well established in *C. elegans* [52]. For example, the NPR-1 circuit oxygen sensing circuit can suppress innate immune responses in nonneuronal tissues [53]. Neuronal INS-7 non-cell-autonomously suppresses intestinal immunity through DAF-2/insulin signaling during infection [54], and GABAergic enteric neurons promote PMK-1-dependent epithelial immunity through FLP-6 neuropeptide [55]. Beyond immune regulation, the *Providencia* JUb39 (an *Enterobacteriaceae*) can bypass sensing circuits through direct production of tyramine that alters sensory behavior of the host [56]. Similarly, chemosensory detection of *P. aeruginosa* phenazines can activate DAF-7/TGF-β neuroendocrine signaling in ASJ neurons to promote pathogen avoidance [57]. Nuclear receptor NHR-86 detects *P. aeruginosa*-derived pyocyanin to activate immune defenses, and *ahr-1*-deficient animals are more susceptible to pyocyanin-overproducing strains [58], establishing that nuclear receptors serve as metabolite-sensing pattern recognition receptors in *C. elegans*.

Our findings extend this landscape of neuronal microbe sensing from acute pathogen responses and behavioral modulation to continuous regulation of commensal community composition.

Parallel host genetic pathways contribute to microbiome composition through distinct mechanisms: DAF-2/16 insulin signaling controls specific taxa including *Ochrobactrum*, and its disruption also results in altered responses to pathogenic *Stenotrophomonas* strains [24,59], while DBL-1/BMP signaling acts through secreted intestinal effectors including lysozymes and C-type lectins to regulate Enterobacteria [60]. The AHR-1 to FLP-8 pathway extends this logic from single-pathogen threat detection to continuous monitoring of commensal community metabolic output. *C. elegans* encodes 274 nuclear hormone receptors, many in chemosensory neurons, and distributed metabolite sensing by this family may broadly coordinate host physiology with microbiome composition beyond what is currently characterized. The apparent transcriptional independence of PMK-1/p38 [61], DAF-2/DAF-16, and SKN-1 in *ahr-1* mutants suggests that this neuroendocrine axis operates in parallel to canonical immune signaling rather than through it.

Among genes downregulated in *ahr-1* mutants, *arrd-11*/TXNIP stands out as significantly reduced, while upregulated genes include the UDP-glucuronosyltransferases *ugt-6* and *ugt-25*, the copper-binding *cnc-2*, and *nnt-1*/NNT, a mitochondrial transhydrogenase implicated in NADPH-dependent redox balance, collectively suggesting broad disruption of both thioredoxin-based and NAD(P)H-linked antioxidant systems despite intact SKN-1. SKN-1/NRF2 canonically regulates *gst-4* expression and is required for full induction of phase I/II detoxification genes under oxidative stress [62,63]. SKN-1 is transcriptionally intact in *ahr-1* mutants despite elevated GST-4 activity and microbiome dysbiosis, indicating that AHR-1 controls intestinal redox through SKN-1-independent transcriptional mechanisms. This is consistent with prior evidence that AHR-1 and SKN-1 regulate *gst-4* through partially non-overlapping mechanisms specific to endogenous metabolite contexts [47,64]. The upstream kinase MEKK-3 may provide a point of convergence between these pathways, as it regulates AHR-1-dependent dietary restriction responses [65] and cooperatively promotes SKN-1 nuclear localization with NSY-1 under oxidative stress [66]. Beyond redox effectors, *ahr-1* mutants display a distinct metabolic signature including elevated branched-chain amino acids and altered polyunsaturated fatty acid composition [19]. This could modify intestinal nutrient availability for colonizing bacteria, contributing to community shifts through a metabolite-mediated mechanism independent of redox tone. Commensal indoles also extend *C. elegans* healthspan through AHR-1 suppression [67], and AHR-1 activity is protective against bacterially produced tryptophan metabolites but counterproductive under certain xenobiotic exposures [21]. This positions the AHR-1 to redox to microbiome feedback as part of a broader circuit in which the metabolic output of the gut community tunes host physiology through the same receptor that controls microbiome composition.

The dysbiosis of *ahr-1* mutants, with *Enterobacterales* and *Pseudomonadales* expansion and *Sphingobacteriales* depletion, mirrors patterns documented in human inflammatory bowel disease, Parkinson disease, and metabolic syndrome [68–70]. AHR pathway interactions with longevity networks are conserved across *C. elegans*, mice, and humans [71], and AHR displays antagonistic pleiotropy in aging [72]. The molecular effectors we identify show conservation in human intestinal epithelium. Human intestinal organoids treated with the AHR ligand FICZ upregulate TXNIP and ARRDC2 (alpha-arrestin orthologs of *arrd-11*) and downregulate GSTA1 and GSTM2 (NRF2-regulated GST orthologs of *gst-4*) [73], mirroring what we observe in *C. elegans*. TXNIP and ARRDC2 belong to the alpha-arrestin family, which can inhibit thioredoxin reductase activity to elevate oxidative stress [74]. The coordinate AHR-dependent regulation of both thioredoxin- and glutathione-based systems points to broad redox remodeling as a conserved output. *C. elegans* GST-4 is alpha-class with high glutathione peroxidase activity, while intestinally expressed mammalian orthologs are predominantly mu-class (GSTM) with lower peroxidase activity, yet both are regulated by AHR in the same direction, suggesting functional convergence despite class divergence. In mammals, microbial indole derivatives protect against colitis through AHR-dependent barrier mechanisms [75], and urolithin A activates AHR-NRF2 to upregulate tight junction proteins [76], paralleling the AHR-1/SKN-1 co-regulation of gut homeostasis identified here. Disrupted AHR signaling through genetic variation, environmental toxin exposure, or dietary ligand deficiency could shift microbiome composition toward stress-tolerant taxa through dysregulated GST and alpha-arrestin/TXNIP expression. The nearly complete rescue of *ahr-1* mutant dysbiosis through intestinal *gst-4* knockdown suggests that interventions targeting downstream effectors rather than the receptor itself may restore community composition even when upstream signaling is impaired.

AHR-1 represents just one of the many ways that hosts monitor microbiome metabolic outputs via expansive repertoires of nuclear hormone receptors. The unique feature here is the specific and targeted response that regulates a specific functional group of bacteria. Thus, metabolic sensing based feedback circuits could represent ancient forms of host-microbiome homeostasis in metazoans.

## MATERIALS AND METHODS

### Maintenance of *C. elegans*

Strains of *C. elegans* used in this study were maintained on *Escherichia coli* OP50 at 20°C for at least three generations prior to conducting experiments. Some strains were provided by the CGC, such as N2 Bristol, DV3805, JV2, MGH171, NY2078, PT501, and ZG24. N2 background strain PHX5588 [syb5588 [*ahr-1*::wrmScarlet::AID::3xFlag]] was constructed by and obtained from SunyBiotech Co., Ltd. All other strains constructed for and used in this study are as follows: BIG0108, BIG0110, BIG0112, BIG0113, BIG0115, BIG0116, BIG0117, BIG0118, BIG0119, BIG0120, BIG0121, BIG0122, BIG0123, BIG0124 [**Table in S2 Table**].

### Bleach synchronization of animals

Animals were grown on *E. coli* OP50 seeded Nematode Growth Media (NGM) plates, and allowed to reach gravid adulthood. Once gravid, animals were washed off NGM using M9 + 0.01% Triton-X and placed into conical tubes. Collected gravid animals were then exposed to a 2:1 mixture of bleach and 5 molar (M) sodium hydroxide to induce cuticle disruption for egg collection. Collected eggs were then rotated overnight in M9 + 0.01% Triton-X to allow for synchronized hatching of axenic, germ-free L1 larvae animals, which were then used for desired assays.

### Preparation of complex community of microbes

LB agar was poured into rectangular Nunc OmniTray Single-Well plates and allowed to solidify overnight. A 96-well plate version of the complex community, BIGbiome [24], was stamped onto the rectangular LB plate using a 96 pin microplate replicator. Bacteria were allowed to grow for two days at 26°C (as a few BIGbiome members take more than 24 hours to sufficiently grow). All colonies of BIGbiome bacterial strains were then cultured in a 2-mL deep 96-well plate using the microplate replicator, and allowed to shake at 26°C for 48 hours covered using a Breathe-Easy membrane. Once grown, optical densities (ODs) were taken using BioTek Cytation 5 and Gen5 software. All bacteria were then combined at a 1:1 ratio after normalizing ODs to 1. This mixture was then spun down for 20 minutes at 4,000 RPM, and subsequently resuspended in equal volume of M9. Combined bacteria were then seeded onto 6 mm NGM plates, and allowed to dry and grow overnight at room temperature before being used for assays. This community consists of microbes outlined in **S1 Table**.

### RNA sequencing sample preparation and analysis

#### Sample preparation and sequencing

N2 Bristol wildtype and ZG24 [*ahr-1* (ia03)] animals were grown on *E. coli* OP50. After a minimum of three generations, gravid animals grown on *E. coli* OP50 were subject to aforementioned bleaching to allow for synchronous populations. Germ-free L1 larvae from the bleaching of each strain were placed onto 90 cm NGM plates that were seeded with either the control *E. coli* OP50 or the experimental BIGbiome complex community. After three days on the bacterial lawns, Day 2 adult animals were transferred to corresponding new lawns away from progeny, using a 40 µM Nylon mesh filter, to avoid starvation until collection at Day 3 of adulthood. Day 3 adults, totaling 120 hours of exposure to each feeding condition, were once again filtered with a 40 µM Nylon mesh filter to ensure a homogeneous population of Day 3 adults. Samples for N2 and ZG24 on OP50 and BIGbiome were prepared and collected in tandem. Roughly 300 animals of Day 3 adults of both strains in both microbial conditions were placed into microcentrifuge tubes, resulting in 12 samples to create the following experimental groups (N2-OP50, ZG24-OP50, N2-BIGbiome, ZG24-BIGbiome). Once collected into 1.5 mL microcentrifuge tubes, 1.25 mL of Trizol was added to a 250 µL pellet of *C. elegans* in M9. These samples were then subject to manual RNA extraction by freezing via liquid nitrogen, then transferred to and thawed in a 37°C water bath. This process was repeated 5 times to complete the mechanical lysis, followed by vortexing and resting at room temperature, repeated 5 times. 100 µL of chloroform was then added, samples were hand-shaken for 15 seconds, then incubated at room temperature for 3 minutes. After incubation, samples were centrifuged at 13,000 rpm at 4°C for 15 minutes. The aqueous phase was transferred to a new, RNAase free 1.5 mL centrifuge tube, where 250 µL of isopropanol was added. The aqueous phase and isopropanol were gently mixed, then incubated for 10 minutes at room temperature. After incubation, the mixture was centrifuged at 13,000 rpm at 4°C for 15 minutes. The supernatant was removed, and the remaining pellet was washed with 500 µL of 75% ethanol, vortexed, and centrifuged at 13,000 rpm at 4°C for 5 minutes. The supernatant of the washed pellet was then removed, and the pellet was placed under an RNAase-free hood to dry for 20 minutes. Once dried, the pellet was dissolved in 50 µL of RNAase-free water (adapted from [77]). The quality and concentration of the extracted RNA was assessed using Agilent 4200 TapeStation, Qubit 4.0 Fluorometer, and DeNovix DS-11 Spectrophotometer Nanodrop. Samples were then submitted to GeneWiz for library preparation and Illumina HiSeq sequencing.

#### Processing and initial analyses of sequencing results

Quality control of RNA sequencing results was performed using FastQC [78]. Reads were then trimmed using Terminal command “bbduk” from the “bbmap” package [79]. Resulting trimmed reads were then mapped to the reference *C. elegans* genome PRJNA13758 version WS283 [39] using STAR aligner [80] with approximately 20 million unique reads per trimmed sample. Trimmed and mapped reads were then processed using an in-house RStudio script using DESeq2 [81,82] that provided the following for transcripts: a tibble table, a raw counts table, and individual Excel files comparing the base mean, log_2_ fold-change (log_2_FC), raw p-values, and adjusted p-values, which were calculated using the Benjamin-Hochberg method. The same trimmed reads were also run through a similar in-house R script using DESeq2 to summarize the transcripts into genes, having transcripts of the same gene summed to properly express DEGs in place of isoforms. RNA-seq based visualizations were also made in RStudio, using outputs from the DESeq2 scripts.

#### Enrichment analyses

Gene sets were extracted from the complex venn and were then subject to analyses by using the Gene Set Enrichment (GSEA) tool on wormbase.org [39]. Genes in subsets were queried with a q-value significance cutoff of 0.1. Resulting GO terms were then plotted as a horizontal bar graph, with the x-axis representing the enrichment fold change of genes in the GO term, the y-axis representing the GO term, and the bar color signifying q-value. Cross-dataset enrichment analyses were completed using datasets published on WormExp v2.0 [83].

### Knockdown of host genes by RNA interference (RNAi)

#### Animal preparation

Animals were fed and reared on OP50 *E. coli* for at least three generations prior to use. Gravid adults were bleach synchronized as mentioned above. Germ-free L1 larvae were placed onto NGM plates seeded with RNAi clones of interest to induce knockdown of the gene of interest.

#### Bacterial preparation

Desired RNAi clones were streaked freshly from the -80°C freezer onto LB agar plates containing 25 µg/mL carbenicillin, and allowed to grow overnight at 37°C. Resulting bacterial colonies were then placed into liquid LB containing 25 µg/mL ampicillin, and grown overnight shaking at 37°C. Overnight cultures were spun down, concentrated 10X, and seeded onto RNAi NGM 12-well plates containing 1 mM IPTG and 50 ug/mL carbenicillin.

#### Experimental procedure

Germ-free L1 larvae were placed onto RNAi NGM 12-well plates onto seeded RNAi clones of interest. At L4, these animals were then transferred to lawns of BIGbiome in a 12-well plate format, and allowed to feed on BIGbiome until collection at Day 3 of adulthood. At Day 2 of adulthood, animals were transferred to new lawns of BIGbiome if needed, away from progeny, to avoid starvation prior to collection day.

### Auxin-induced protein degradation (AID) assays

6 cm NGM plates were either seeded with a final concentration of 4 millimolar (mM) naphthaleneacetic acid (K-NAA) and allowed to dry, or contained a final concentration of 4 mM K-NAA prior to pouring, to induce AID - mediated protein knockdown. The plates were then seeded with the desired bacterial strain(s) or community mixture after allowing plates to sit overnight.

Animals were sync-hatched overnight in M9 to L1 larvae, and grown until L4 on NGM without K-NAA, then placed onto NGM plates either containing or lacking (control condition) 4 mM K-NAA seeded with desired microbial lawn.

### Fluorescent reporter data acquisition and analyses

Transcriptional reporter and biosensor images were captured using a Nikon TiE inverted epifluorescent scope, quantified using ImageJ [84], with the collected quantification later processed in R. For *flp-8*p::GFP reporter (NY2078) assays, images were excited at 488 nanometers (nm) with a 2 millisecond (ms) exposure time at 40X magnification. In resulting images, neurons were outlined and the total signal found in the outlined areas were used as output fluorescence values for analyses. For *gst-4*::GFP reporter assays, images were captured using an automated 96-well plate imaging setup at 4X. Animals were excited at 488 nm with a 30 ms exposure time. Plug-in WormAlign (see Supplemental Information), was used to trace, straighten, and capture average fluorescence across animal length for the desired animals using pixel count and normalization by worm length. This output was then used in plug-in PixelProcessor to assign fluorescence values based on the pixel matrix data obtained from WormAlign. Outputs from PixelProcessor were then used as input data for RStudio visualizations using an in-house script. For JV2 biosensor strain quantifications, animals were excited at both 395 nm (Grx) and 470 nm (roGFP) with an exposure time of 40 ms and emission captured at 535 nm at 20X magnification. Animals were imaged in two portions, and images were processed using an in-house script that obtained an image-based GSSH/2GSH ratio by dividing the fluorescence from Grx by fluorescence found in roGFP (adapted from [85]). Output processed images were then stitched together in ImageJ with a “Fire” Look-up Table (LUT). Processed images from RNAi conditions were observed at a max of 1.3 and stitched linearly, compared to no RNAi conditions at a max of 1 and stitched using max blending.

### Bacterial H_2_O_2_ exposure assays

Desired bacteria were grown shaking overnight at 26°C. 200 µL of overnight cultures was plated into a flat-bottom 96- well plate, and exposed to various final concentrations of 3% hydrogen peroxide (H_2_O_2_) ranging from 0 millimolar (mM) to 20 mM (0 mM, 500 µM, 1 mM, 2.5 mM, 5 mM, 10 mM, 20 mM). Culture growths following exposures were monitored in triplicate by OD600 in a BioTek Cytation 5 plate reader every 30 minutes over the course of 24 hours. The area under the curve after 24 hours was obtained via an in-house R script, and subsequently plotted.

### Microbiome sequencing analyses

Sequences were obtained from a core facility that were demultiplexed. Demultiplexed samples went through processing methods employing the use of QIIME 2 following a modified protocol of the “Moving Pictures Tutorial”. Analyses were conducted using QIIME 2 version 2020.2 – 2023.2 [86] standard protocol. Quality control was performed with the QIIME 2 quality-filter q-score plugin. Error correction and denoising were carried out using Deblur [87] was used to perform error correction and denoising using a fixed trim length of 120 bp. The resulting feature table was summarized with QIIME 2 feature-table summarize to provide sample sequencing depth information. Representative sequences were tabulated using QIIME 2 feature-table tabulate-seqs for downstream taxonomic and phylogenetic analyses. Diversity analyses were performed using the core-metrics-phylogenetic pipeline in QIIME 2 with a rarefaction depth of 1,103 sequences per sample. Both alpha diversity and beta diversity metrics were calculated. Feature tables were exported from QIIME 2 for downstream analysis and converted to tab-delimited text format using BIOM v2.1.5 [88].

Representative sequences were exported in FASTA format via the QIIME 2 visualization interface. Representative sequences were taxonomically annotated using NCBI BLAST+ [89], having a custom nucleotide BLAST database that was constructed from the BIGbiome reference collection.

Sequences were queried against this database with blastn, using a stringent alignment threshold of ≥99% identity and 100% query coverage. Hits were then used for downstream taxonomic assignment. All outputs of QIIME 2 and downstream taxonomic assignment were then used as input data for visualization and statistical analyses using ATIMA [90] and in-house RStudio scripts.

### Microbial gene enrichment analyses

Microbial functional enrichment analyses were performed through Integrated Microbial Genomes and Microbiomes (IMG/M) extension from Joint Genome Institute (JGI) [91]. Mann-Whitney statistical tests were performed on microbes grouped by whether they were increased or decreased in an *ahr-1* mutant or knockdown microbiome. These results detailed which KEGG Ontology (KO) pathways and enzymes were statistically significantly different between the two groupings. Genomic comparison analysis visualizations were performed using input data used to create WormBiome [28] in conjunction with RStudio, while utilizing the information from the functional KEGG Ontology analyses as a guide.

### Figure illustrations

Artificial intelligence (AI) ChatGPT (model 4) [92] was used to draft and troubleshoot RStudio scripts used for illustrations. Some illustrations were also created with BioRender [93].

## ACKNOWLEDGMENTS

We would like to thank additional members of the Samuel lab (Dana Blackburn, Audrey Parish) for providing feedback on experiments, data analyses, and data visualizations. Additionally, we would like to thank Dr. David Reiner, Dr. Anne-Marie Krachler, Dr. Robert Britton, Dr. Roy Sillitoe, and Dr. Sundararajah Thevananther for providing support and feedback on the project at critical stages.

Thank you to the Center for Metagenomics and Microbiome Research sequencing center at Baylor College of Medicine for performing microbiome sequencing and assisting with initial processing. We would also like to thank the *Caenorhabditis* Genetics Center (CGC) – which is funded by NIH Office of Research Infrastructure Programs (P40 OD010440) – for providing transgenic and mutant *C. elegans* worm strains.

## SUPPORTING INFORMATION AND DATA AVAILABILITY

All data is available within the manuscript and Supporting Information files. Numerical and raw data datasheets used to create graphical visuals can be found in **S1 Data** (**Figs 1-7**). Uncropped images can be found in **S1 Raw Images** (**Figs 3-5**). Raw Z-stack images are deposited on Zenodo (Fig 3, https://doi.org/10.5281/zenodo.20672162). Microbiome metadata can be found in **S2 Data** (Fig 1, Fig 3, and Fig 6), as microbiome sequencing is deposited on Zenodo (https://doi.org/10.5281/zenodo.20603022). RNA-sequencing raw data and supplemental file (count table) have been deposited in NCBI Gene Expression Omnibus (GEO) under accession number GSE310883.

**S1 Fig: Broader view of AHR-1 dependent transcriptional networks highlights alterations in immune and stress response pathways.** Extended analyses of Fig 2, including differentially expressed genes (DEGs) in *ahr-1* mutants exposed to OP50 and/or BIGbiome compared to respective N2 (wildtype) controls. **(A)** Heatmap of DEGs upregulated on both OP50 AND BIGbiome. Genes are grouped by functional categories, and 35 uncharacterized genes were not plotted. **(B-C)** Summary of significant cross-dataset enrichments *ahr-1*-dependent DEGs in either OP50 OR BIGbiome and published datasets (False Discovery Rate (FDR) <0.05). Barplots detail the count of genes that were upregulated, downregulated, or genetic targets (N/A; e.g., transcription factor (TF) binding datasets) in published datasets (WormExp). Bubble size represents gene counts that overlap between datasets and *ahr-1*-dependent DEGs. Categories of published datasets are noted, including some with combined impacts (e.g., pathogen and environmental stress).

**S2 Fig: Microbiome-dependent transcriptional responses in *ahr-1* and wildtype animals.** Extended analyses of Fig 2, focusing on microbiome-based responses that are altered or retained in *ahr-1* mutant or N2 (wildtype) animals. **(A)** Scatterplot displaying log_2_ fold change (log2FC) differences in *ahr-1* mutant and/or N2 animals on BIGbiome versus *E. coli* OP50. Differentially expressed genes (DEGs) that maintain significant responses to BIGbiome compared to OP50 (*ahr-1* independent) are plotted in grey (1,320), while DEGs that respond only in N2 (687) or *ahr-1* (1,014) are colored blue and red, respectively. **(B)** GO analyses of microbiome-dependent DEGs in each of the conditions are plotted. Color represents Q-value significance, while dot size represents absolute value of enrichment fold change. **(C-E)** Summary of cross-dataset enrichments based on microbiome-dependent DEGs in N2 and/or *ahr-1* mutants with published datasets (False Discovery Rate (FDR) <0.05). Barplots detail genes sum of upregulated, downregulated, or genetic targets (N/A; e.g., transcription factor (TF) binding datasets) in datasets (WormExp). Bubble size represents gene count overlap between datasets and microbiome-dependent DEGs. Categories of published datasets are noted, including some with combined impacts (e.g., pathogen and immune).

**S3 Fig**: **Microbiome-dependent alterations in FLP-8 reporter expression in *ahr-1* mutants.** In addition to changes in expression of *flp-8*p::GFP in URX neurons (see Fig 3), we also observe microbiome-dependent changes to *flp-8*p::GFP expression in other neurons. Boxplots of fluorescence quantification of expression in **(A)** AUA head interneurons and **(B)** PVM mid-body mechanosensory neurons is presented for N2 (wildtype) and *ahr-1* mutants on either *E. coli* OP50 or BIGbiome. Significance was determined using a Wilcoxon test (N2 vs. *ahr-1* mutants; **p<0.01,***p<0.001). **(C)** Representative images of *flp-8*p::GFP expression in PVM mid-body neurons (small grey arrows) and ectopic expression of *flp-8*::GFP in unidentified tail sensory neurons (open white arrowheads) on OP50 in *ahr-1* mutants. Scale bar is 100 µm.

**S4 Fig: Adulthood knockdown of AHR-1 increases redox tone while decreasing oxidative stress responses in a BIGbiome-dependent manner.** Similar to Fig 4, assessment of redox tone and general oxidative stress responses was performed in AHR-1::AID animals. **(A-B)** Fluorescence quantification and representative scaled images of redox tone (oxidization level based on plasma color scale lookup table (LUT)). Scale bar is 250 µm. **(C-D)** Fluorescence quantification and summed scaled images (of all animals in condition) of *gst-4*p::GFP reporter strains. Significance is noted in compact letter display (CLD) format (samples with shared letters = no significant difference); Kruskal-Wallis statistical tests followed by Dunn-Bonferroni post-hoc tests for comparison of fluorescence in different genetic backgrounds for each set of mutants. Raw data used to create graphical visualizations can be found in **S1 Data**.

**S5 Fig: Intestinal knockdown of downstream regulators of AHR-1 have minimal impact on gut microbiome of wildtype animals.** Similar to Fig 6, intestinal-specific RNAi was completed for N2 (wildtype) and *ahr-1(ia03)* mutants. **(A)** PCoA plots of between sample beta diversity metrics (weighted Bray-Curtis dissimilarity) faceted by host genetic background. Box plots of RNAi knockdowns in *ahr-1* mutants that contribute to normalization of gut microbiome metrics toward N2 empty vector (EV, L4440) controls, including: **(B)** weighted Bray-Curtis distances, **(C)** within sample alpha diversity (Shannon Index), and relative abundance of gut bacteria orders **(D)** and strains **(E)** that bloom in *ahr-1* mutants. Kruskal-Wallis test followed by Dunn-Bonferroni post-hoc test for comparison of significance for relative abundance of each order between RNAi conditions represented by compact letter format calculated using microbial order and strain facets. Significance is displayed in the compact letter format.

S6 Fig: Genomic profiles of oxidative stress response genes in microbes that exhibit altered gut colonization of *ahr-1* mutant animals. Gene counts for a given KEGG Ontology (KO) term identified within the genome of a given bacterial strain are plotted based on size and color. Box color behind taxonomic groups highlight consistent colonization profiles of bacterial orders: grey, no change or mixed; red, higher in *ahr-1* mutants; blue, lower in *ahr-1* mutants.

**S1 Table: List of BIGbiome microbial strains.**

**S2 Table: List of *C. elegans* strains used in this study.**

**S1 Data: Spreadsheet containing the raw data used for all plotted graphs in the manuscript.**

**S2 Data: Spreadsheet containing metadata files for all microbiome sequencing datasets. S1 Raw Images: Uncropped flattened images presented in this manuscript.**

